# Machine Learning Guided Optimization of an Oral Microemulsion System: A Bayesian Optimization Approach

**DOI:** 10.64898/2026.01.30.702790

**Authors:** Madeline Gunawardena, Bao Chau, Hayden Nothacker, Rudra Pangeni, Thomas Roper, Qingguo Xu, Charles McGill

**Affiliations:** Department of Pharmaceutics, School of Pharmacy, Virginia Commonwealth University, Richmond, VA 23298, USA; Department of Chemical and Life Science Engineering, College of Engineering, Virginia Commonwealth University, Richmond, VA 23219, USA; Department of Ophthalmology, Department of Pediatrics, Department of Biomedical Engineering, Center for Pharmaceutical Engineering, Center for Drug Discovery, Massey Cancer Center, Virginia Commonwealth University, Richmond, VA 23298, USA

**Keywords:** Oral drug delivery, drug formulation development, property prediction, physicochemical regression, stability classification

## Abstract

Oral microemulsions are one drug delivery system often implemented to improve intestinal permeability and oral bioavailability of poorly-water soluble drugs. They also present several practical advantages including high patient compliance and simplified manufacturing methods which contribute to their promise as marketable drug products. Despite these advantages, however, the microemulsion formulation development process is extremely time consuming and resource intensive, typically involving extensive screening of components and excessive preliminary trials in order to achieve stable formulations. As a result, MEs are often suboptimal in their therapeutic performance, and there is an unmet need for improved methods to streamline microemulsion formulation design. In this work, a batch Bayesian Optimization strategy was used to design a subset of unique in-specification microemulsions with highly optimized physicochemical properties in five iterations containing batches of five experiments. A two-phase modeling approach was developed to achieve this goal and allowed for navigating a complex experimental design space, including multiple oils, surfactants, cosurfactants, and processing parameters with a training dataset consisting of 22 experiments. As a result of this study, five high-performing blank microemulsions were identified, and four of them exhibited physical stability upon storage for up to 30 days. Further, when loaded with two different model drug candidates, three of the microemulsions achieved high drug loading, acceptable stability and improved in-vitro permeability, highlighting their promise for potential translation to later stages of development.

## 1. Introduction

Oral microemulsions (MEs) are thermodynamically stable drug delivery systems that employ oil, surfactant/cosurfactant, and aqueous phases to solubilize a drug compound of interest. Compared to alternative delivery systems, oral MEs offer several advantages related to both therapeutic performance and manufacturing of the dosage form, some of which include increased drug solubility and permeability in the intestinal tract, improved patient compliance, enhanced storage stability, and simplified manufacturing methods(1,2). For these reasons, use of the ME delivery system is desirable for applications in which these aspects stand to be improved.

Despite their many therapeutic advantages, the formulation development of oral MEs can be quite inefficient(2), ultimately limiting their utility and scalability as potential drug products. Determining a stable composition that meets predefined therapeutic and manufacturing requirements often involves heuristic trial and error approaches(2,3). In these cases, formulation scientists will select a small subset of oils and surfactants to test against one another and iteratively vary their ratios until a stable ME is achieved. Given a particular set of components, this thermodynamically stable experimental space, or “microemulsion region” as it is often named, is extremely narrow compared to the full experimental space. It is exceptionally challenging and time consuming to design a high-performing, optimized composition, as the experimental operating space is highly constrained and only a limited number of combinations can be practically tested(2,3).

To adopt a more methodical approach, many recent studies have employed quality by design (QbD) principles. Design of experiments (DoE) is one tool commonly used to implement QbD, which allows for assessing the impact of different input factors on output responses(4,5). In a ME context, input factors may include different material attributes and processing parameters, which ultimately impact measurable performance-related outcomes of the formulation(4,5). Through recent work, our team developed a high-performing, first-generation oral ME utilizing Box-Behnken Design (BBD) - one type of DoE which is often employed for its efficiency and applicability to process optimization. With all other factors held constant, the BBD campaign evaluated different percentages of oil, ratios of surfactant to cosurfactant, and sonication times, with the goal of achieving desirable physicochemical properties that would enhance oral delivery of a novel PPARα agonist A190(6). The first-generation formulation designed in this work was able to achieve desirable physicochemical properties and drug content within the design space, and further in-vitro and in-vivo characterizations revealed its promise as a potential oral formulation(6).

Despite these outcomes, however, there are several key limitations associated with the DoE approach that should be considered. Primarily, the oils and surfactants used in the optimization campaign needed to be determined before experimentation to reduce the number of parameters and decrease the required number of experiments(7). This excluded the possibility of testing other, potentially high-performing candidates. In this case, only Oleic acid, Tween 80, and PEG 400 were evaluated, rather than the larger pool of ME excipients reported and used in literature. Secondly, all compositions needed to be fixed before testing, which left no room for guided sampling or iterative optimization processes(8). Other notable limitations included limited parameter resolution and poor sampling efficiency. Considering these disadvantages, it is clear that oral MEs stand to be further improved through the application of more advanced methods, which can improve upon these weaknesses and provide more clinically relevant results. Algorithmic optimization techniques that leverage machine learning modeling can be one potential tool to remove the constraints seen in DoE, while optimizing and exploring a larger design space with greater efficiency(9).

Bayesian optimization (BO), an iterative optimization algorithm that enables the efficient optimization of black-box functions, is traditionally applied in machine learning problems to find the global optimum for expensive-to-evaluate functions (e.g. hyperparameter tuning)(10–13). Recent innovations have led to the adoption of BO in medicinal chemistry, process chemistry, and formulation optimization(8,10,14). As evidenced by these innovations, using this technique enables the exploration of larger design spaces without the scaling issue typically seen in DoEs(7,11). BO is designed to balance exploration of the design space with exploitation of the current known optimal conditions(11,13,14). Compared to other optimization methods, BO can effectively navigate a more complex design space through the iterative sampling and model retraining cycle to consistently obtain high-performing configurations in fewer experiments(8,10). In this application, BO’s efficiency in complex optimization problems allows for an increase in dimensionality of the previous ME design space, removing the limitation of the DoE optimization campaign by including additional categorical parameters (additional oils, surfactant, and cosurfactant) and continuous working ranges rather than discrete values for continuous parameters (oil volume, surfactant ratio, and sonication time).

Traditional BO only suggests a single configuration to test in each iteration cycle, prohibiting the ability to conduct experiments in parallel. In practice, parallel experimentation tends to be the most efficient method for conducting experiments in terms of time and resources(15). Recent work with batch BO methodologies, multiple experimental configurations are suggested, removing the previous limitation and further increasing the throughput and efficiency of the optimization campaign(13,16,17).

In this study, we created a batch BO algorithm that allowed us to methodically and efficiently identify a series of blank ME compositions with highly optimized physicochemical characteristics. We proposed that using this approach would allow us to significantly expand the experimental design space beyond what was enabled by the DoE campaign and introduce a novel platform that can be used to design multiple high-performing MEs. To test this hypothesis, we designed the algorithm using a two-phased approach and used it to prompt the formulation of 25 compositionally-diverse MEs. Further, to evaluate the translatability of the best MEs, the top 5-performers were further tested via loading with two model drugs, A190 and Fenofibrate, and characterized to confirm physicochemical performance, stability upon short-term storage, and desired in-vitro permeability.

## 2. Materials and Methods

### 2.1 Materials

Capryol 90 (propylene glycol caprylate), Maisine CC (glyceryl monolineoleate), Labrafil M 1944 CS (oleoyl polyoxyl-6 glycerides), and Labrasol (caprylocaproyl polyoxyl-8 glycerides) were gifted by Gattefossé (Saint-Priest, France). Propylene Glycol (1,2-propanediol), Transcutol HP (diethylene glycol monoethyl ether), Cremophor RH 40 (castor oil, ethyloxylated), Tween 80 (polyethylene glycol sorbitan monooleate), Oleic Acid (cis-9-octadecenoic acid), PEG 400 (polyethylene glycol 400), and Fenofibrate were purchased from Sigma-Aldrich (MO, USA). A190 (99% purity) was provided by Dr. Adam Duerfeldt from University of Minnesota. Parallel artificial membrane permeability assay (PAMPA) plates were purchased from Thermofisher Scientific (Waltham, MA, USA).

### 2.2 Selection of Microemulsion Components

Many different biocompatible oils, surfactants and cosurfactants have been utilized in preclinical oral ME formulations. As a result, there are many which could be reasonably considered in this Bayesian optimization campaign. However, in order to define a feasible and relevant experimental space, a discrete subset of these components needed to be selected. A preliminary list of several candidate oils, surfactants, and cosurfactants was first identified, then components on the list were screened against one another in a preliminary miscibility evaluation. Those which exhibited poor miscibility with at least two candidates of a different component class were eliminated. For example, if a given oil was immiscible with two or more surfactants, it was removed from the list. Results from this miscibility evaluation can be seen in the **supplementary information (Tables S2-S4)**.

### 2.3 Preparation and Physicochemical Characterization of Microemulsion Formulations

#### 2.3.1 Microemulsion Formulation

All MEs evaluated in this work were formulated using a standardized protocol to maintain consistency between samples. Briefly, MEs were formulated using the high energy ultrasonication method(18), which involved adding the oil, surfactant, and cosurfactant phases in series with 90 seconds of mixing time in between each addition (Benchmark BV1010, USA) and a 30 minute water bath sonication (Branson CPX2800H, USA) after all phases had been added. Further sonication via an ultrasonic Liquid Processor VCX 500 (Sonics & Materials, Inc., CT, USA) was used in order to ensure homogeneity within each sample, with the total duration treated as one of the input factors. If any of the formulations showed visual instability after sonication (i.e., clear phase separation, drug precipitation, or excessive turbidity), they were classified as “failed” and not pursued for further evaluations.

Three replicates of each ME (n=3), each at a total volume of 1 mL, were prepared and evaluated to account for any procedural variability associated with the process.

#### 2.3.2 Physicochemical Characterization of Microemulsions

Mean droplet size, polydispersity index (PDI), and zeta potential of each formulation were measured after 24-hour storage at room temperature using a dynamic laser light scattering analyzer (Malvern Zetasizer Nano ZS90; Malvern Instruments, Malvern, UK). To prepare the samples for such analyses, they were diluted 1:100 v/v with 10 mM NaCl solution and sonicated for 1 minute via the Branson Ultrasonics Bath (Branson CPX2800H, USA) to ensure appropriate dispersion and minimize multiple scattering effects. Measurements were carried out at 25 °C, and three separate runs were recorded for each physicochemical parameter for each ME replicate.

### 2.4 Bayesian Optimization Pipeline Development

#### 2.4.1 Prior Dataset

In order to begin the optimization campaign, a screening set of experiments consisting of 5 previous optimum formulations and 10 quasi-randomized formulations was used. All experiments fell within the relevant design space described in **Table 1**. The formulations within the screening set were pulled from the team’s prior work(6), and the quasi-randomized experiments were constructed using a randomizer script with constraints that encouraged an even distribution of formulations, thereby further exploring the preexisting design space. More specifically, the randomizer script implemented the following constraints: (i) each oil, surfactant, and cosurfactant appeared in at least two of the ten formulations; (ii) at least three of the ten formulations had a surfactant ratio above or below 1:1; and (iii) all oil, surfactant, and cosurfactant combinations were compatible with one another according to the predefined miscibility matrix (**Tables S2-S4 in SI**). All formulations were created in triplicate, visually assessed for phase separation, and physicochemically characterized according to the previously described methods.

**Table 1.** Design space of blank microemulsion optimization campaign. Parameters and respective settings or ranges involved in the optimization campaign. Note that PEG 400 and Tween 80 were available to be used as both surfactant and cosurfactant candidates. This decision was based upon prior formulation experience as well as prevalence of use in related literature studies.

| Parameter | Design Space |
| --- | --- |
| Oil Volume (%) | 7.5 - 22.5 |
| Surfactant:Cosurfactant Ratio (Smix) | 3:1 - 1:3 |
| Sonication Time (min) | 0 - 3 |
| Oil | Oleic Acid, Capryol 90, Soybean Oil, Maisine Oil, Capmul MCM |
| Surfactant | PEG 400, Tween 80, Tween 20, Labrasol |
| Cosurfactant | Tween 80, Transcutol HP, Propylene Glycol, Ethanol, PEG 400 |

#### 2.4.2 Categorical Descriptors

To provide supplementary information about the categorical parameters within the design space (i.e., oils, surfactants, cosurfactants), descriptors were included as additional input parameters to the models. Hydrophile-lipophile balance (HLB) values for each component were used to inform the surrogate models on their degree of hydrophilicity or lipophilicity(19). Binary measures of compatibility between each of the candidate ME components, as determined in the miscibility evaluation, were also used as descriptors. See **Table S1 in SI** for a more detailed overview.

#### 2.4.3 Dataset Preprocessing

To further preprocess the screening dataset, all continuous parameters were normalized to a range of 0 to 1 using MinMax scalers. To simplify the way in which the surfactant composition was handled during modeling, the ratios shown in **Table 1** were instead represented as equivalents of surfactant. For any given formulation, there were 12 equivalents total, and there could be anywhere from 3 to 9 equivalents of surfactant, with the remainder being cosurfactant. All categorical parameters were preprocessed utilizing one-hot encoding and used to assign categorical descriptors. The descriptors shown in **Table S1 in SI** were attached to the categorical parameters with HLB values assigned to each normalized oil, surfactant, and cosurfactant. Compatibility values were implemented directly as their raw values, as compatibility was initially expressed as either 0 or 1 responses and thus, did not require further preprocessing.

The compatibility values were derived from three secondary data tables, Oil–Surfactant Compatibility (**Table S2**), Oil–Cosurfactant Compatibility (**Table S3**), and Surfactant–Cosurfactant Compatibility (**Table S4**), which are provided in the **SI**. The compatibility, along with normalized HLB values for each component, supplied additional chemical context beyond the primary formulation variables of interest. During dataset construction, the pipeline loaded each table and assigned every formulation a set of descriptor values derived from the specific oil, surfactant, and cosurfactant present. Descriptor columns were then appended directly to the encoded model inputs and used in every stage of the workflow, including the Phase 1 classifier, the Phase 2 regression models, and the Bayesian optimization routines. The goal here was to provide the models with supplementary quantitative information surrounding component compatibility and interaction patterns, which helped to shape how the models interpreted the formulation landscape and influenced the scoring and selection of new candidate formulations during optimization.

#### 2.4.4 Objective Function Determination

A minimization objective function was also created for use during modeling to represent all ME outputs as a single value. Outputs were the same as those considered in the original DoE campaign(6), along with one additional parameter, phase separation. The outputs considered are listed in **Table 2**, along with their respective goals and target criteria for success.

**Table 2.**
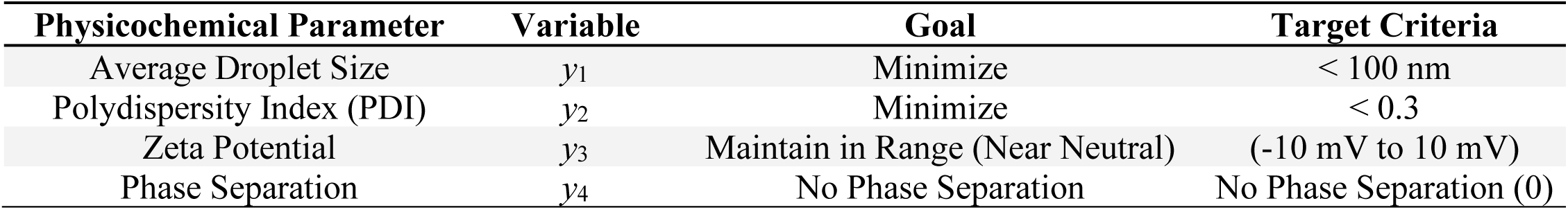
Response targets for formulation optimization.

**Equation 1** presents the objective function (J) used, which combines all the individual output parameters (*y*) with their respective weights (*w*). A weight of 1 was assigned to all response targets, except for phase separation, which was assigned a value of 10 to prioritize the selection of stable formulations prior to optimizing performance. **Equations 2-4** were used to normalize the specific output parameter between 0 and 1 and implement the target criteria into the objective function. **Equations 2 and 4** represent droplet size (*y_1_*) and zeta potential(*y_3_*), and their functions are equal to zero when the statement in the Iverson bracket is false. This indicates that once the target criteria are met, further improvement in either criterion provides no significant additional benefit. The opposite is true for **Equation 3**, where PDI (*y_2_*) has a decrease in influence by ¼ after reaching the target criteria. Weighting and objective function decisions were made based upon previous formulation experience as well as experimental goals. Phase separation (*y*_4_) was bounded to the range of [0,1] in the dataset and machine learning classification predictions with 0 indicating no phase separation (stable) and 1 if displaying any phase separation (unstable).

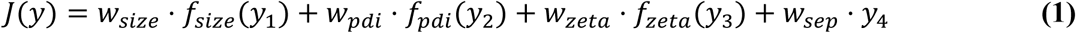

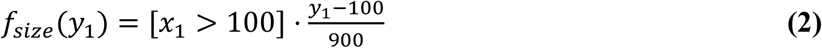

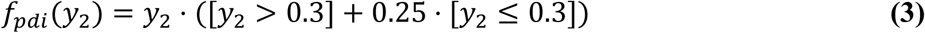

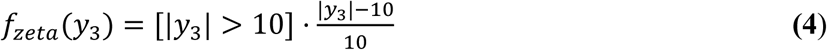

2.4.5 *Surrogate Model Development*

To support model-guided exploration of the ME formulation space, a two-phase surrogate modeling framework was developed. In Phase 1, a probabilistic classifier was used to estimate the likelihood that a candidate ME would remain physically stable (i.e., not phase separate). In Phase 2, uncertainty-aware regression models were used to predict physicochemical outcomes (mean droplet size, polydispersity index (PDI), and zeta potential) for said ME, *conditional on predicted stability*, thereby focusing the optimizer on feasible regions of the design space. This architecture follows constrained BO strategies that pair a feasibility model with uncertainty-aware surrogates to avoid sampling unproductive or infeasible regions(20). **Figure S1 in SI** presents a visual representation of this workflow.

#### 2.4.6 Phase 1: Stability Classification

During the first phase, a classification model was trained using all 54 experimental formulations using formulation using formulation phase-separation as the target. Two classifier types were evaluated for this model - a Random Forest (RF) classifier and a Multilayer Perceptron (MLP) classifier. Because feasibility probabilities were used directly within the constraint component of the acquisition function, probability calibration was treated as an essential metric for surrogate model selection. Both models employed post-hoc probability calibration within a nested five-fold stratified cross-validation workflow(21). Outer folds produced unbiased out-of-fold (OOF) probabilities for evaluation and plotting (**Figure S2 in SI**), while inner folds tuned hyperparameters using negative log-likelihood (NLL) as the primary objective. All reported probability-based metrics and diagnostic plots were computed strictly from OOF predictions (**Table S5 in SI**).

#### 2.4.7 Phase 2: Physicochemical Regression

In ME design and optimization studies, parameters such as droplet size, polydispersity index, and zeta potential can only be treated as relevant quality attributes for stable, single-phase systems. Measurements resulting from unstable or phase separated systems are not meaningful for guiding formulation development(22). With this in mind, Phase 2 surrogate development was designed to use only the subset of formulations that were observed to be physically stable (n = 39) in Phase 1. Candidate model surrogates included Gaussian Process Regression (GPR) and Multilayer Perceptron Ensembles (MLP ensembles), both of which provide predictive means and uncertainty estimates required for uncertainty-aware BO(23). Regression benchmarking used nested five-fold cross-validation. Outer folds produced OOF predictions used to compute RMSE, MAE, R², and NLL, while inner folds tuned hyperparameters. Because BO depends critically on usable uncertainty estimates(24), NLL and calibration served as the primary probabilistic criterion, with RMSE and R² used as secondary indicators of mean prediction accuracy when likelihood performance was similar. Final Phase 2 surrogate selection and diagnostic interpretation are addressed in the results and discussion section. Based on the combined performance metrics and uncertainty diagnostics, the final Phase 2 surrogate set consisted of GPR models for droplet size and PDI, and an MLP ensemble model for zeta potential, integrated downstream of the Phase 1 RF feasibility classifier for constrained BO.

#### 2.4.8 Batching Method Development

A hallucinated liar method was implemented in order to guide the batching of experiments, rather than the traditional one experiment per iteration. This method is reliant on simulating the suggested configuration (hallucinating) and assigning poor performance for the response targets (liar/lying) to encourage batch diversity by punishing suggested configurations in the cycle(12,25). More specifically, the batching method began via the standard BO workflow with a suggested configuration (Experiment 1), and the response target values were assigned the default phase-separated formulation values (droplet size = 10,000, PDI = 1, zeta potential = 100, Phase Separation = 1). The configuration and response target values were then appended to the dataset, creating a new temporary dataset which could be pushed back into the BO cycle to obtain another suggested configuration. The cycle ended when the number of suggested configurations equaled the designated batch size. See **Figures S1 and S5 in SI** for visual representations of this workflow.

#### 2.4.9 Meshing Method Development

A meshing method was created to evaluate the objective function (J(*y*)) and calculate the acquisition function (expected improvement) at mesh points across the design space. The meshing first considered the categorical parameters, creating all possible combinations in a list and assigning their designated categorical descriptors. Each combination was then used to form a design space hypercube (coarse mesh) created using only the continuous parameters. Hypercubes were created by evenly spacing out the normalized continuous parameters with a mesh size dictating the number of levels between the upper and lower boundaries. All categorical combinations and their respective hypercubes were sampled at all points and analyzed using the surrogate models to obtain the predicted mean and uncertainty. Categorical combinations were then sorted by the highest acquisition function value seen in their hypercube. The top 10% of categorical combinations were carried forward to fine continuous meshes. For each of the combinations’ fine mesh, a narrower hypercube was centered around the best point in the coarse mesh. If the fine mesh passed the design space boundaries, the extra points were removed and not made available for analysis. All fine mesh points were analyzed in the same manner as the coarse mesh. The highest acquisition function value was selected as the best experiment for a given iteration.

### 2.5 Iterative Collection and Selection of High Performing Microemulsions

During batching, MEs were formulated in five separate batches of five each (25 total). Upon completion of a given batch, physicochemical data was measured, reported as an average value, and added back into the dataset. The meshing method was used once again to identify the next set of promising compositions to be tested. After all five batches were completed, weighted objective scores were calculated for each completed ME **(Equation 1**) to evaluate outcomes. An objective score closer to zero indicated higher performance and closer alignment with the target criteria. The five formulations showing the lowest objective scores were identified as the highest performing MEs and selected for further assessment.

### 2.6 Drug Loading and Secondary Assessment of Candidate Formulations

#### 2.6.1 Reformulation, Drug Loading, and Short-Term Stability Assessment of Microemulsions

Following the optimization campaign and selection of the top MEs, all five were reformulated at a total volume of 5 mL. In addition to the blank ME form, each composition was also loaded with A190 or Fenofibrate, which served as model drugs, to evaluate potential translatability to drug-loaded form. Although the campaign did not involve drug inclusion, it was important to evaluate if the MEs could successfully encapsulate a drug of interest. Target loadings were set at 0.5 % (w/v) across all formulations for consistency. Briefly, the appropriate amount of A190 or Fenofibrate was first dissolved in the oil phase and sonicated for 30 minutes to ensure complete dissolution. All sequential formulation steps were the same as those previously described. Following formulation, the MEs were stored at room temperature for 24 hours prior to analysis. To mirror the standard protocol used during modeling, any formulation which showed instability upon visual inspection (phase separation, drug recrystallization, or excessive turbidity) was classified as “failed” and was not evaluated further. Stable MEs were followed for 30 days to assess short-term shelf life and drug product stability, with physicochemical characterization and drug content assessments occurring on days 1, 7, 15, and 30.

#### 2.6.2 Drug Content Determination

Drug content was determined using high-performance liquid chromatography methods (HPLC-UV, Shimadzu Prominence LC System). Samples were diluted with acetonitrile, filtered using 0.22 um membrane filters (PTFE, hydrophobic, ThermoScientific), and injected into an Agilent Pursuit XRs 5 C18 column (250 x 4.6 mm) with UV-detector system. For both A190 and Fenofibrate, mobile phases consisted of acetonitrile and water, each containing 0.1% trifluoroacetic acid at a flow rate of 1mL/min, and ratios of 70:30 v/v and 10:90 v/v aqueous:organic, respectively. Injection volumes remained constant at 10 uL, and peak detection was performed at 227 and 284 nm, respectively. This procedure was repeated for each stability timepoint.

#### 2.6.3 Evaluation of Microemulsion In Vitro Permeabilities

To further assess the promise of the model-resultant MEs and gain insight into their intestinal permeabilities, an in-vitro parallel artificial membrane permeability assay (PAMPA) was performed for the stable drug-loaded MEs. Donor samples consisted of either the free drugs (A190 or Fenofibrate in PBS), free drug-dispersions (A190 or Fenofibrate dispersed in a mixture of 10% PEG 400 + 5% DMSO + 85% of 10% HP-β CD), or the drug-loaded MEs. 0.3 mL of PBS (pH 6.8) was loaded into each well of the acceptor plate, and acceptor and donor plates were sandwiched, ensuring that the membrane of the donor plate was in contact with the media in the acceptor plate. 0.2 mL of each sample was then loaded into the appropriate donor well, and the entire plate assembly was incubated for 5 hours at room temperature. Following incubation, samples were collected from the acceptor and donor plates, and drug concentrations of all samples were measured via previously described methods. Effective permeability of each sample was quantified using **Equation 5**.

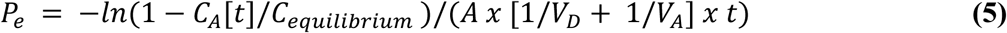

where *P*_e_ is the permeability (cm/s), *A* is the effective filter area (0.228 cm^2^), *V*D is the volume of the donor well (0.2 mL), *V*_A_ is the volume of the receptor well (0.3 mL), *t* is the total time of incubation in seconds, *C*_A_(*t*) denotes the concentration of drug in the receptor well at time *t*, and *C*_equilibrium_ represents (*C*_D_[*t*] × *V*_D_ + *C*_A_[*t*] × *V*_A_)/(*V*_D_ + *V*_A_). *C*_D_(*t*) denotes the concentration of drug in the donor well at time t.

#### 2.6.4 Statistical Analysis

All statistical analyses were performed using GraphPad Prism 10.1 (GraphPad Software Inc., San Diego, CA, USA). Student’s t-test was used for comparing two experimental groups, and one-way or two-way ANOVAs were used for comparing three or more groups. Differences were considered statistically different if p < 0.05.

## 3. Results and Discussion

### 3.1. Phase 1 Surrogate Model Comparison and Selection (Classification)

To enforce feasibility constraints during BO, the Phase 1 classifier must provide reliable and well-calibrated estimates of formulation instability. Two candidate models, a MLP and a RF, were evaluated for their ability to separate physically stable and unstable formulations while producing interpretable probability estimates suitable for use within a constrained acquisition function.

While both models were able to achieve strong class discrimination, their probability structures differed substantially. The MLP produced smoother, intermediate probability outputs that resulted in ambiguous feasibility estimates for some stable formulations, whereas the RF generated sharply separated, low-entropy probability assignments that more closely aligned with the observed binary separation behavior. Diagnostic analyses including probability distributions, calibration curves, and confusion matrices are provided in **Figure S3 in SI**. These analyses demonstrate that the RF classifier yielded better-calibrated and more decisive feasibility probabilities, minimizing the risk of falsely rejecting viable formulations during optimization. Accordingly, the calibrated RF classifier was selected as the Phase 1 surrogate model for all subsequent constrained Bayesian optimization analyses.

### 3.2. Phase 2 Surrogate Model Comparison and Selection (Regression)

Phase 2 surrogate modeling was performed exclusively on the subset of 39 formulations that remained physically stable following Phase 1 screening. Regression surrogates were trained to predict droplet size, polydispersity index (PDI), and zeta potential using nested five-fold cross-validation. Model evaluation emphasized not only point-estimate accuracy, but also the quality of predictive uncertainty, as BO directly exploits uncertainty through the acquisition function.

Across all targets, candidate models exhibited target-dependent performance, necessitating separate surrogate selection for each physicochemical response. Quantitative cross-validated performance metrics and probabilistic diagnostics are summarized in **Table S6 and Figure S4 in SI**. While multiple models were able to achieve competitive accuracy under metric-based screening, uncertainty calibration behavior proved decisive for final selection.

Based on the combined evaluation of predictive accuracy, likelihood performance, and uncertainty calibration, GPR models were selected for droplet size and PDI prediction, while a MLP ensemble was selected for zeta potential. This surrogate set provided the most reliable balance of accurate mean predictions and usable uncertainty estimates for integration into the BO framework.

#### 3.2.5 Progressive Surrogate Model Stability Under Iterative Data Acquisition

To assess the robustness of the Phase 2 surrogate models under iterative BO, a progressive retraining analysis was performed in which models were updated after each experimentally executed batch of formulations recommended by the optimization framework. Model performance was evaluated using a fixed holdout test set of 13 physically stable formulations (Phase Sep = 0) drawn exclusively from the initial 66-experiment dataset, selected once via randomized sampling with a fixed seed and retained for all future evaluations. This design isolates the effect of incremental incorporation of newly generated experimental results from successive BO batches on surrogate predictive performance while eliminating confounding effects due to changes in test-set composition.

Following execution of each optimization batch, Phase 2 surrogate models were retrained using cumulatively expanded training datasets: the initial dataset alone, followed by the initial dataset plus Batches A through E. For each iteration, the same surrogate architectures and training procedures were used, consisting of GPR models for droplet size and PDI, and a MLP for zeta potential as described previously.

Surrogate performance on the fixed holdout test set is summarized in **Figure 1** (Stability of Phase-2 Surrogate Model Performance on Iterative Batches), with RMSE and R² shown for droplet size (panels **A–B**), PDI (panels **C–D**), and zeta potential (panels **E–F**) as functions of progressive training-set expansion. Across all three physicochemical targets, predictive performance remained largely stable as the total number of experiments increased from 66 to 141. Despite more than a two-fold increase in training data from the initial campaign to Batch E, droplet size and PDI surrogates exhibited no systematic loss of accuracy, with only modest, non-monotonic fluctuations across intermediate iterations.

**Figure 1.**
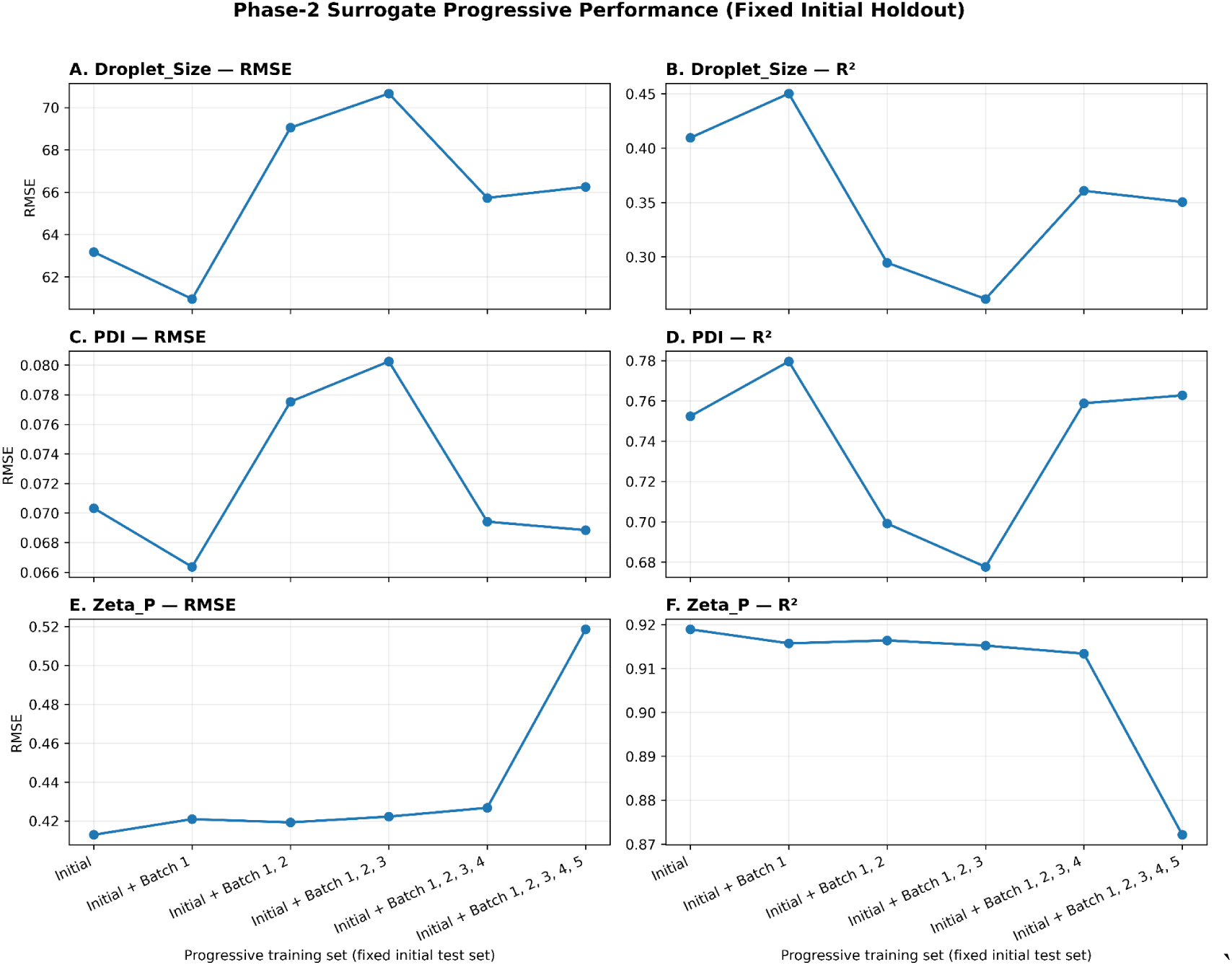
Stability of Phase-2 Surrogate Model Performance on Iterative Batches.

For zeta potential, surrogate performance remained effectively unchanged through Batches A–D before exhibiting a modest decline upon inclusion of Batch E, with R² decreasing from approximately 0.91 to 0.87 and RMSE increasing from approximately 0.42 to 0.52 (**Figure 1E–F**). This represents the largest performance change at Batch E, whereas droplet size and PDI recouped their generalization metrics at this time. The magnitude of the zeta potential shift remains limited relative to the overall prediction scale and does not indicate catastrophic loss of generalization. Overall, results indicate that Phase 2 surrogate performance remains robust under progressive incorporation of BO-recommended experimental batches.

### 3.3 Batch Bayesian Optimization Microemulsion Formulation Optimization Campaign

Following completion of the five batches and tabulation of the resultant physicochemical data, all formulations, including those in the starting datasets and those formulated during iterative learning, were scored via the weighted objective score described previously. The five formulations which showed the lowest objective scores and thus were selected for further investigation are shown below in **Table 5**, with their physicochemical outcomes shown in **Table S7 in SI**.

**Table 5.** Top-Performing Microemulsions from Bayesian Optimization Campaign.

| Formulation | Objective Score | Oil | Oil % (v/v) | Surfactant | Cosurfactant | Smix | Sonication Time (min) |
| --- | --- | --- | --- | --- | --- | --- | --- |
| B4 | 0.0424 | Capryol 90 | 8.250 | Tween 20 | Ethanol | 1.70:2.30 | 1.650 |
| D3 | 0.0464 | Capryol 90 | 7.875 | Tween 80 | Tween 80 | 1.05:2.95 | 0.825 |
| D5 | 0.1098 | Maisine Oil | 7.500 | Tween 80 | Propylene Glycol | 1.25:2.75 | 1.725 |
| F5 | 0.1208 | Capryol 90 | 10.000 | Tween 20 | PEG 400 | 2.00:2.00 | 2.000 |
| E2 | 0.1343 | Capryol 90 | 8.325 | Tween 20 | Transcutol HP | 1.85:2.15 | 2.025 |

With the selection of the surrogate models for each of the response variables, the two-phase multimodal batch BO pipeline began the optimization campaign, displayed in **Figure 2**, for the ME formulation blanks. In the final dataset, there were 13 formulations grouped at an objective score of 31, indicating phase separation within 24 hours of formulation.

**Figure 2.**
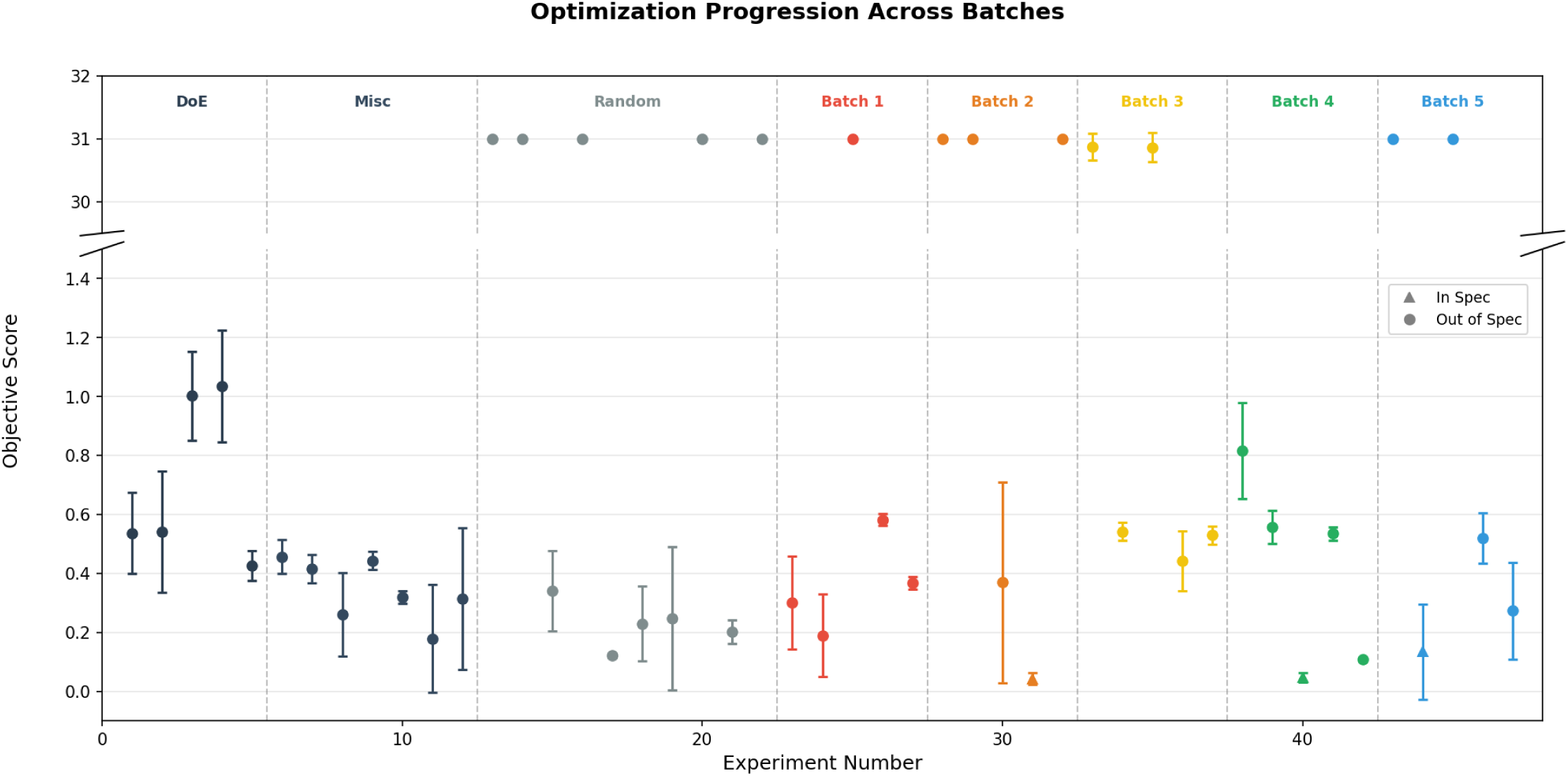
Optimization Progression Plot of the Optimization Campaign.

Despite the high percentage of failed formulations, the BO method discovered three in-specification formulations (B4, D3, and E2) in five iterations of batches of five compared to the zero from the DoE campaign. The combination of a large quantity of failed formulations and low in-specification formulations highlights the complexity of the design space and the narrow microemulsion region. The addition of formulations with high variability in stability (Eg. B4, Random 7) created a challenge for surrogate model accuracy because of the complex nature of semi-stable formulations.

Visualization of the optimization campaign displaying the results of the batches compared to the initial dataset, comparing the previous optimized design of experiment formulations (DoE), miscellaneous formulations (Misc), and randomized with constraints (Random) to the optimization batches. Triangle markers indicate the formulations that fell within specifications.

Despite these challenges, the optimization campaign discovered nine formulations that perform better on average than the previous optimal formulations revealed through DoE (DoEOPT). Seven out of the nine formulations utilized Capryol 90, which was identified early as a high performing oil and led to the three in-specification MEs discovered through optimization in surfactant, cosurfactant combinations and continuous parameter optimization. Due to the low iteration count, the amount of fine-tuning optimization for continuous parameters was minimal and did not converge at a singular value for the Capryol 90 formulations besides oil volume, which converged near the lower boundary condition. After the high-performing MEs were discovered, BO started to exploit Capryol 90 formulations which is why it remained the most popular oil in each iterated batch. This bias can be explained by the included miscellaneous data, which contained only Capryol 90, Tween 80 and Transcutol P formulations and displayed strong but out-of-specification performance. Had those formulations been more compositionally diverse, it is likely that the resultant formulations would mirror that. Compared to DoEOPT, all three in-specification formulations performed between 3.2 to 10 times better, with B4 and D3 having smaller batch-to-batch deviation. Within the optimization campaign, convergence was achieved with oil and oil volume set at Capryol 90 and the lower oil range (7.5-10.0), respectively.

### 3.4 Interpretation of Selected MEs and Short-term Stability Evaluations

As previously described, the target criteria for each of the physicochemical parameters were as follows: < 100nm, < 0.3, and between -10 and +10 mV for average droplet size, PDI, and zeta potential values, respectively. Among the 5 MEs, droplet sizes ranged from 11.69 ± 1.05 to 148.14 ± 2.11 nm, PDIs from 0.170 ± 0.081 to 0.296 ± 0.001, and zeta potentials from -4.88 ± 1.46 to -2.23 ± 0.24 mV. Notably, three of the five formulations were able to meet or exceed all three target criteria (B4, D3, and E2). Additionally, D3 contained a formulation with the same surfactant and cosurfactant selected which resulted in the Smix value becoming irrelevant as all 12 equivalents are assigned to Tween 80. The two which failed to meet the criteria, D5 and F5, showed higher droplet sizes than desired (>100nm). However, they did still meet the target criteria for PDI and zeta potential (**Table S7 in SI**). Though these droplet sizes were larger than desired for this optimization campaign, they did still fall within the acceptable droplet size range defined for oral MEs(26), and thus, may still present promise as candidate formulations. Despite this result, the subset of formulations investigated were high performing and exhibited good trending toward the desired target criteria.

Based on the magnitude of the available experimental space in this campaign (**Table 1**), reaching this endpoint with a relatively small and fixed experimental cap (i.e., only 25 formulations total) is highly promising. Compared to standard trial-and-error based methods typically employed for ME formulation, this approach presents a much more efficient development platform(7,27). Considering the same experimental space, identifying three successful formulations through trial-and-error would have used exponentially more time and resources, causing significant process delays which would ultimately hinder translation of the MEs to later stages of drug development(2,3).

Beyond discovery of the optimized compositions themselves, another major benefit of the batched BO approach is its ability to identify formulation patterns and provide further insight into how excipient selection and processing parameters may affect experimental outcomes(27,28). With this in mind, there are some important compositional trends to be noted. Primarily, Capryol 90 was used as the oil component in four of the five top-performing formulations, and the tween surfactants (Tweens 20 and 80) in all five formulations. This suggests that compared to the other candidate excipients in the design space, Capryol 90, Tween 20, and Tween 80 may present advantages when it comes to physicochemical properties. However, due the small experimental cap of 25 experiments, it should also be recognized that the full experimental space, spanning all possible excipients and combinations, could not be explored. Additionally, the compositional range that was explored within the initial screening set represents only a small fraction of this total experimental space, so it is also very likely that the algorithm’s prompting of formulations using these excipients was based on that discrete prior knowledge. With these limitations in mind, it is clear that many other successful regions of the design space remain unexplored and thus unknown. Additionally, oil volumes for all formulations remained at the lower end of the possible range (< nearly 10% v/v in a possible range of 7.5-22.5%) and sonication times tended to sit around the middle region of the experimental range (nearly 1.5 or 2 minutes in a possible range of 0-3 minutes). The exact range at which each of these parameters is optimal cannot be determined, as it is highly dependent on the excipients at hand. However, these trends can still be meaningful in understanding the experimental space that was explored.

Interestingly, Smix ratios across the formulations revealed that there are patterns of higher levels of cosurfactant versus surfactant. As reported in prior literature, ME systems tend to contain the opposite (more surfactant than cosurfactant), as the role of the cosurfactant is to enhance and supplement the role of the primary surfactant(29). However, this is not always the case, and in this campaign, the cosurfactants are seemingly more important in driving physicochemical properties to the desired level than are primary surfactants. It is also important to highlight that the surfactants and cosurfactants used in this study were initially classified as either one or the other prior to modeling. However, in practicality, a particular surfactant is not limited to one or the other. Depending on the system at hand, it can serve as both - so the surfactant and cosurfactant roles identified here may be working oppositely in some cases.

In preparation for short-term stability evaluations, all five top-performing MEs were reformulated at 5mL volume. Following 24 hours of storage at room temperature, D3 showed clear phase separation, and thus, was not evaluated further. The other four formulations, however, remained visually transparent and showed no signs of phase separation. Images of each formulation can be seen in **Figure S6 in SI,** and physicochemical results at the initial timepoint as compared to the initial formulations created at a 1mL volume are shown in **Table S8 in SI**. The four formulations were followed upon storage for up to 30 days as intended, and property trending over time can be seen in **Figure 3**.

**Figure 3.**
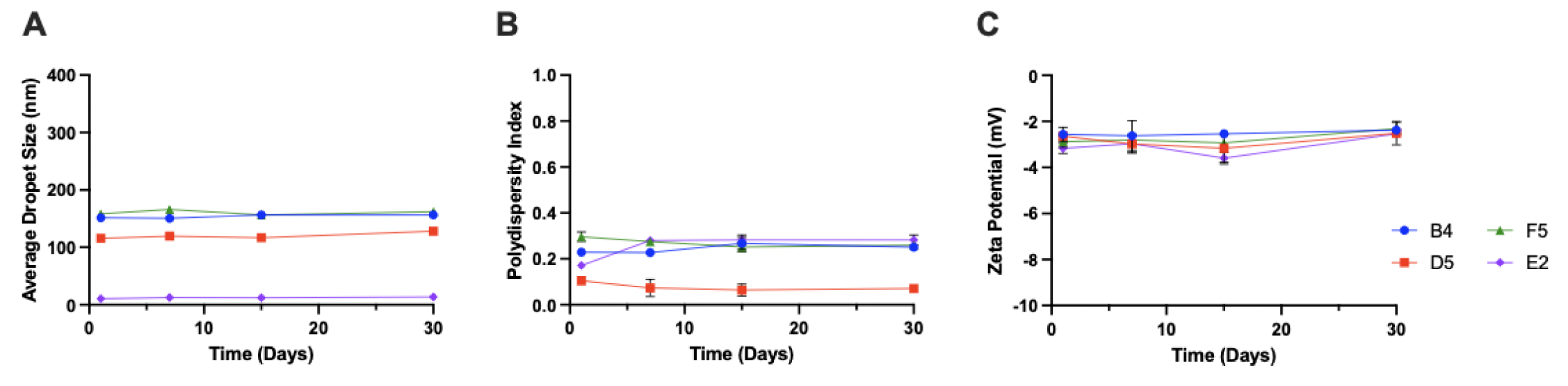
Physical stability of model-optimized microemulsions during 30 days of storage at room temperature. **(A)** Average droplet size, **(B)** polydispersity index, and **(C)** zeta potentials are shown for the four blank MEs which passed the visual stability inspection.

One surprising result of the process, however, is that when reformulated at the 5mL volume for the stability study, several of the resulting physicochemical values were different from those observed during the design campaign at 1mL volume (**Table S8 in SI**). This may indicate potential issues with translation of the formulation process from the 1mL to 5 mL scale, specifically relating to the probe sonication aspect. In this case, the amplitude or intensity of the sonication was kept constant in both cases to adhere to the standard formulation procedure. However, the energy per unit volume was changed as a result, meaning that the level of sonication was different between the two different volumes(30). Ideally, volumes used in the stability study would have remained constant with those in the optimization campaign (1mL for both). However, in order to create an adequate amount of formulation in preparation for the stability and drug loading assessments during the 30-day time period, practical limitations led to the five-fold increase in volume. Future work should investigate how to successfully translate the process to different volume scales, even larger than those explored in this work. However, at the 5mL volume, physicochemical properties were still trending toward the desired targets, so in this case, this limitation did not significantly hinder the experimental outcomes.

Overall, the investigated MEs remained physically stable for the evaluation period. Though formulations B4, D5 and F5 had average droplet sizes slightly exceeding the target criteria (<100 nm), values showed minimal fluctuation over time. Polydispersity indexes were also relatively consistent and for all formulations, remained within the target criteria (<0.3) for all timepoints. However, it should be noted that formulation E2 showed an increase in PDI from day 1 to day 7, which may suggest slow equilibration or potentially, early signs of instability. Further, zeta potentials remained in range with minimal fluctuations over time. Although the optimization campaign did not consider or optimize for storage stability as a key outcome, the formulations identified did exhibit stability on the short-term time frame, highlighting the potential of the platform developed to design shelf-stable drug products relevant for future translation to the market. Had the formulations shown instability, their promise would be extremely limited.

### 3.5 Drug Loading and In-Vitro Evaluation of Microemulsions

The optimization campaign revealed important trends within the experimental design space. However, it did not include drug-loaded MEs, rather, it involved the blank ME vehicles themselves. Though it is well known that the drug plays a critical role in a ME’s stability and heavily impacts physicochemical outcomes(31), drug loading was excluded from this campaign because the primary goal was to design a novel optimization platform which can be used as a baseline in future ME formulation development. Had one specific drug compound been considered and utilized throughout this work, the model created would be limited to identifying formulations for only that particular drug, or potentially, other drugs with similar chemical structures and properties. By optimizing the blank systems first, this study identified not only a series of MEs, but also a computational tool that can be extended to a wide range of drugs across different pharmaceutical applications. In support of this goal, however, further investigation into whether the blank MEs identified could successfully load a drug of interest was needed. Specifically, the aim was to understand if the physicochemical properties would be changed in the presence of the drug, as well as if the ME vehicles could achieve high levels of drug loading and short-term physical stability.

To this end, we selected two surrogate drugs of interest, Fenofibrate and A190, which are non-opioid PPARα agonists. Although Fenofibrate and A190 show high potency and isotype selectivity for PPARα compared to alternative agonists, both compounds suffer from poor aqueous solubility and permeability, ultimately leading to suboptimal bioavailability when administered orally (34). Further details regarding their chemical structures and properties can be seen in **Table S9 in SI**. Leveraging the oral ME delivery system, we recognized the potential to significantly improve upon these limitations. This, in combination with previous work from our team(6), makes these two drugs relevant for evaluating our model-resultant formulations. To explore this avenue, the five MEs were also reformulated in drug-loaded form, either containing A190 or Fenofibrate.

When loaded with drugs, the MEs showed similar stability trends, but overall success tended to depend on the specific compound involved (**Figure 4**). For instance, when loaded with Fenofibrate, B4 showed the desired stability with minimal fluctuations in physicochemical properties. However, when loaded with A190, the outcome was quite different. Increases in average droplet size as well as significant fluctuations in PDI occurred during storage, suggesting that the B4 composition may be suitable for some drug compounds, but not others. Further, this result suggests that the chemical structure and properties of the drug loaded are important factors to consider in future optimization campaigns and formulation development as a whole(31). Similarly, for F5, F5-A190 exhibited good stability, but the F5-Feno did not. For E2, despite clear differences in droplet size magnitude (E2-A190 vs the E2-Feno), the formulations both appeared to have relatively stable sizes over time. PDI trending is similar to that of the blank formulation, showing an increase between day 1 and 7. However, as previously described, that may be a sign of slow equilibration or early instability.

**Figure 4.**
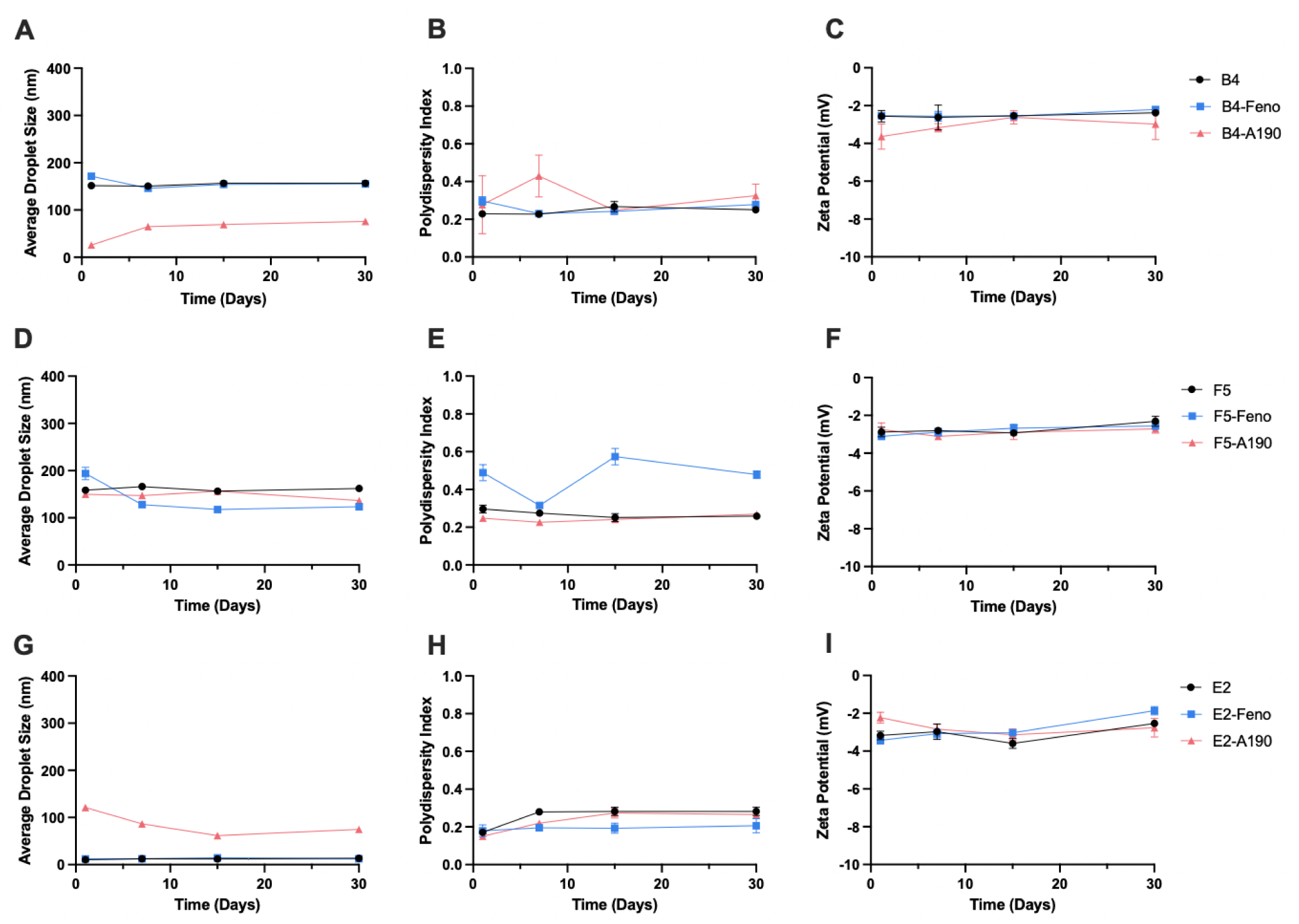
Physical stability of model-optimized microemulsions during 30 days of storage at room temperature. Average droplet size, polydispersity index, and zeta potentials are shown for formulations B4 **(A-C)**, F5 **(D-F)**, and E2 **(G-I)**, each evaluated in the blank, A190-, and Fenofibrate-loaded forms. Feno = Fenofibrate.

When assessed for drug content over the 30 days, all six MEs exhibited a high level of drug content, with all values remaining in the 90 to 110% window of initial drug content for the full 30 days (**Figure S7 in SI**). Similarly, the idea that the formulations can achieve and maintain extremely high drug loading is very impressive considering that the drug was not involved in their initial design. Additionally, the targeted concentration for all three formulations was 0.5 w/v %, which compared to many other formulations on the market or in literature, is relatively high. This value was selected based on previous formulation experience and ultimate clinical goals. However, it is possible that increasing or decreasing the targeted drug loading may have led to different outcomes. The drug, its concentration, and its solubility within a given ME system are all highly important factors for determining stability.

Finally, the six MEs were screened for effective permeabilities using the PAMPA permeability assay (**Figure 5**). Outcomes revealed that for both A190 and Fenofibrate, all three ME compositions showed significantly improved permeabilities compared to the controls (respective free drugs in PBS or dispersions). One of the ME’s major advantages as an oral delivery system is its ability to enhance intestinal permeation in the intestinal tract. This is most often the result of nano-sized droplets with large surface area and the permeation-enhancing effects from nonionic surfactants on the intestinal epithelium(29). In this case, all six MEs significantly increased the permeability of the drug compounds at hand, indicating that the compositions employed would be able to enhance systemic delivery of the drug compound in vivo. In complement with the stability study, these results even further support the success of this optimization campaign. Without considering stability or permeability as responses in this work, we were still able to identify six high-performing MEs which showed successes in these areas.

**Figure 5.**
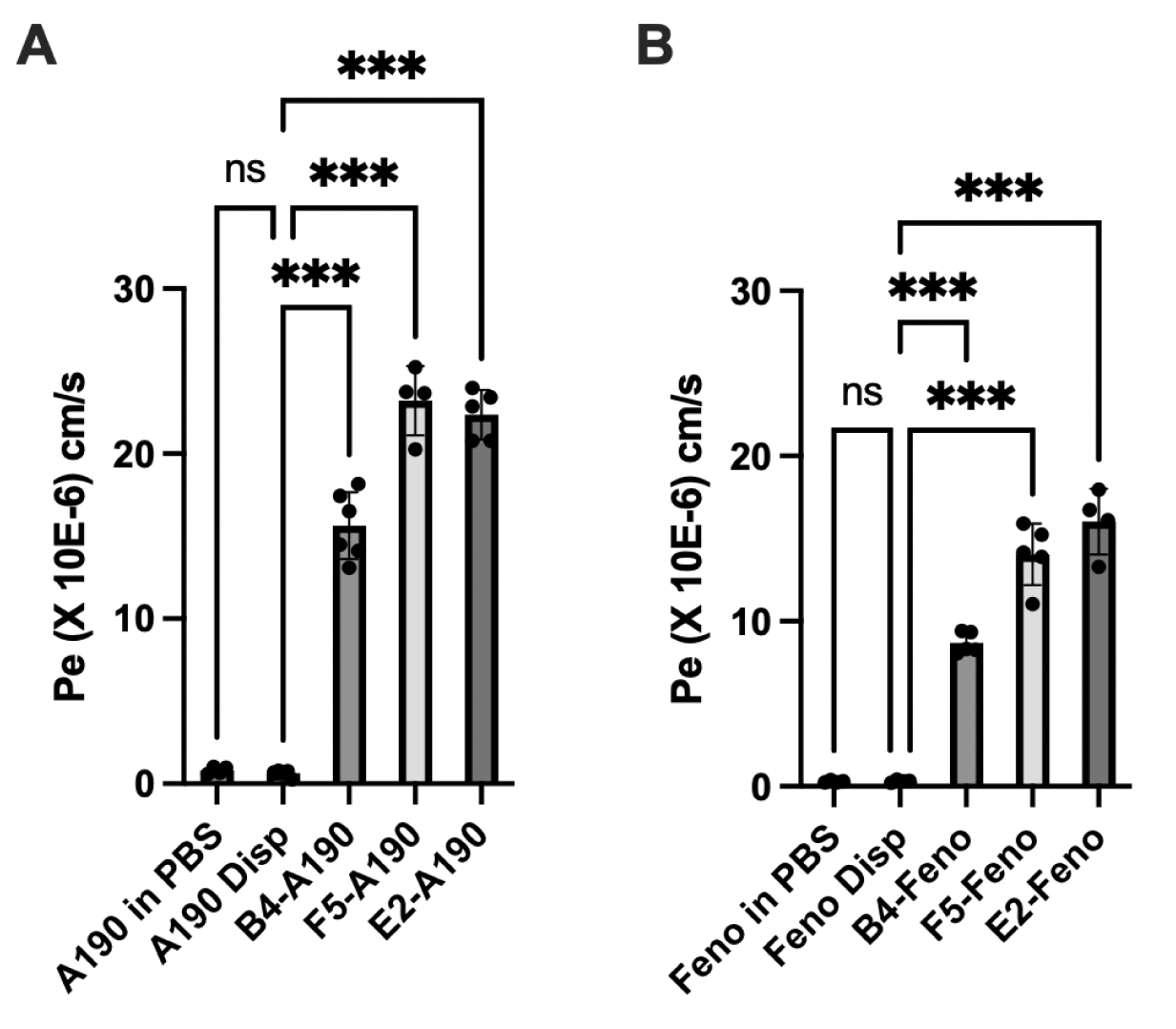
Effective permeabilities of the A190- and Fenofibrate-loaded microemulsions. (**A** and **B**, respectively). Data are represented as Mean ± SD (n=4-6). Disp = dispersion. Feno = Fenofibrate. ns = not significant. *** p < 0.001, compared to the respective drug in dispersion.

In complement with the stability study, these results further support the success of this optimization campaign. Without considering stability or permeability as responses in the modeling side of this work, the team was able to identify 6 high-performing MEs which showed successes in these areas. It should also be noted that the goal of this work was not to determine the “best” formulation or identify significant differences between each composition. Rather, the goal was to identify a set of high-performing formulations and confirm their promise through in-vitro characterization and evaluations. However, for both A190 and Fenofibrate, the F5 and E2 formulations seemingly outperformed B4. They were significantly larger than B4, however, not statistically different from one another (**Table S10 in SI**). This may indicate that those two compositions would achieve better GI permeability and in turn higher bioavailability in vivo, which is the goal associated with using the oral ME system. Interestingly, previous physicochemical outcomes tended to be highly dependent upon the drug at hand, but for permeability, the behavior of the vehicles themselves did not seem to be. Without further investigation of the chemical properties of the drugs themselves, the team does recognize this pattern, and perhaps the success of these two ME compositions could extend to other drug compounds as well, not just for the two modeled here. Future investigations should consider a larger spectrum of drug compounds, specifically those with different properties and structures, to see if outcomes would be different and validate these findings. Overall, these model drugs successfully highlighted the potential for the three ME compositions.

## 4. Conclusion

This study introduced a two-phase batch Bayesian optimization modeling approach for machine learning drug formulation development of oral MEs. Through a small number of experiments, five in-vitro optimized compositions were identified, highlighting the ability of machine-learning based strategies to significantly improve experimental efficiency for ME development. When loaded with Fenofibrate and A190 as model drugs, the compositions also achieved physicochemical stability for 30 days, a high level of drug loading, and significant improvements in vitro permeability compared to the free drug, all of which are hallmarks of the ME system and further support the promise of those identified through this work. This methodology establishes a foundation for shifting away from inefficient trial-and-error based approaches toward more efficient, data-driven development methods. This campaign only explored 25 diverse MEs in order to test and validate the modeling algorithm as a proof of concept. However, future campaigns involving a larger experimental cap, more focused exploration in largely unexplored experimental regions, or even model adaptation to include additional features or target other parameters would be able to further elucidate ME design space. Ultimately, the modeling strategy demonstrated has the potential to streamline ME formulation and extend to other clinically relevant drug compounds and delivery systems, creating a powerful tool for directing formulation development.

## Declarations

### Availability of Data and Materials

All data generated in this work and needed to evaluate the conclusion are presented in the paper or have been made available in supplemental information. Additional data that support the findings of this study are available from the corresponding author upon reasonable request. All code utilized in this work is available on GitHub. [https://github.com/mcgillresearchgroup/Microemulsion_Batched_Bayes]

### Authors’ Contributions

M.G., B.C.: data curation, investigation, methodology, formal analysis, writing-original draft. H.N, R.P: data curation, investigation, methodology, formal analysis, Writing - Review & Editing. T.R., Q.X., and C.M.: conceptualization, investigation, methodology, writing-review and editing, Funding acquisition.

## Acknowledgements

The authors would like to acknowledge Dr. Adam Duerfeldt and Grant Berkbigler at the University Minnesota for providing the A190 that was used in this work as a model drug.

