## Supplementary material for "Machine Learning Guided Optimization of an Oral Microemulsion System: A Bayesian Optimization Approach": Table S1

#### ***General Methods Supplemental Information***

***Table S1.*** *Dataset Descriptors for Handling Categorical Parameters*

| Descriptors | Design Space |
|---|---|
| HLB Value | 1.0 - 16.7 |
| Surfactant - Cosurfactant Compatibility | Yes / No (1/0) |
| Surfactant - Oil Compatibility | Yes / No (1/0) |
| Oil - Cosurfactant Compatibility | Yes / No (1/0) |

***Table S2***. *Oil–Surfactant Compatibility.*

Binary compatibility table identifying feasible oil–surfactant combinations based on prior experimental screening. Entries of 1 indicate compatible pairs retained in the formulation space, while 0 denotes incompatible combinations excluded from modeling and optimization.

| Oil Component | Labrasol | Tween 80 | Tween 20 |
|---|---|---|---|
| Capmul MCM | 1 | 1 | 0 |
| Capryol 90 | 1 | 1 | 1 |
| Maisine Oil | 1 | 1 | 1 |
| Soybean Oil | 1 | 1 | 1 |
| Safflower Oil | 1 | 1 | 1 |
| Oleic Acid | 1 | 1 | 0 |

***Table S3.*** *Oil–Cosurfactant Compatibility.*

Binary compatibility table defining feasible oil–cosurfactant pairings. Compatible combinations (1) are permitted during surrogate model training and Bayesian optimization, whereas incompatible pairs (0) are removed from consideration.

| Oil Component | PEG 400 | Propylene Glycol | Ethanol | Cremophor EL | Glycerin | Transcutol HP |
|---|---|---|---|---|---|---|
| Capmul MCM | 1 | 1 | 1 | 1 | 0 | 1 |
| Capryol 90 | 1 | 1 | 1 | 1 | 1 | 1 |
| Maisine Oil | 1 | 0 | 1 | 1 | 0 | 1 |
| Soybean Oil | 1 | 0 | 1 | 0 | 1 | 1 |
| Safflower Oil | 0 | 0 | 0 | 1 | 1 | 1 |
| Oleic Acid | 1 | 0 | 1 | 1 | 0 | 1 |

***Table S4.*** *Surfactant–Cosurfactant Compatibility.*

Binary compatibility table specifying surfactant–cosurfactant combinations that form stable mixtures under the investigated conditions. This table is applied as a feasibility constraint to restrict the candidate design space during optimization.

| Surfactant Component | PEG 400 | Propylene Glycol | Ethanol | Cremophor EL | Glycerin | Transcutol HP |
|---|---|---|---|---|---|---|
| Labrasol | 1 | 1 | 1 | 1 | 0 | 1 |
| Tween 80 | 0 | 1 | 1 | 1 | 0 | 1 |
| Tween 20 | 1 | 1 | 1 | 1 | 0 | 1 |

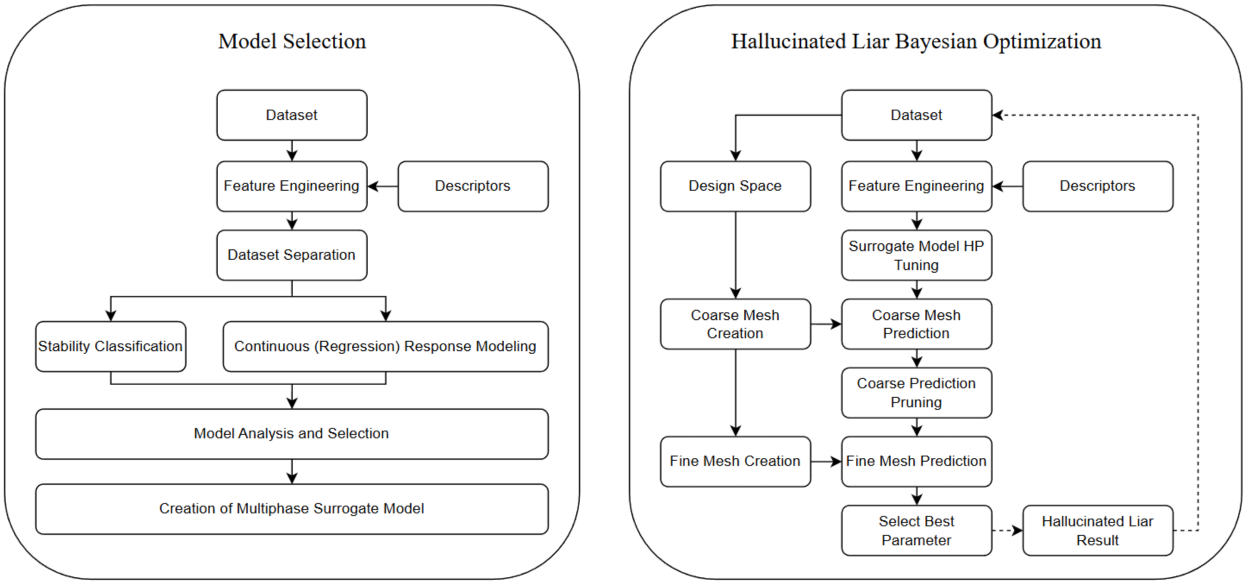


***Figure S1.*** *Surrogate Model Development and Hallucinated Bayesian Optimization Process Flow Diagram* The dataset is preprocessed and appended with categorical descriptors and used to train a multiphase surrogate model consisting of a probabilistic stability classifier (Phase 1) and regression models for physicochemical properties (Phase 2). The multiphase surrogate model evaluates categorical candidates via the coarse categorical mesh, followed by local fine continuous refinement. A hallucinated batch loop iteratively refits the surrogates to propose a batch of five formulations.

#### ***Phase 1 Supplemental Information***

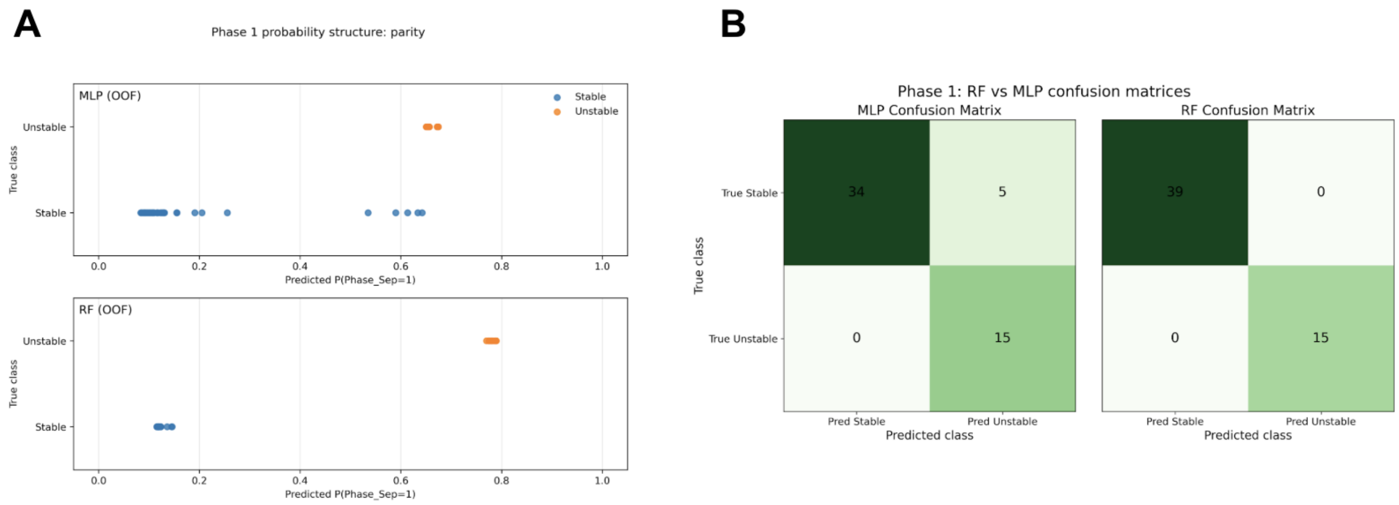


***Figure S2.*** *Phase 1 Modeling Results.* **(A)** Probability parity plots comparing predicted instability probability against true class labels for Phase 1 classification. MLP (top) and RF (bottom) predictions demonstrate class-conditional probability separation, with unstable formulations assigned consistently higher probabilities than stable formulations. **(B)** Confusion matrices summarizing Phase 1 classification performance for MLP (left) and RF (right). Both models achieve perfect identification of unstable formulations, while RF eliminates false positives observed for MLP among stable samples.

##### ***Probability Structure and Class Separation***

To assess the suitability of candidate Phase-1 classifiers for use as feasibility constraints in Bayesian optimization, the probability structures produced by a calibrated multilayer perceptron (MLP) and a calibrated random forest

(RF) were examined in detail. Although both models achieved perfect threshold-independent discrimination (ROC-AUC = 1.00; PR-AUC = 1.00), their predicted probability distributions differed substantially.

The MLP classifier assigned many stable (Class 0; non-separating) formulations intermediate predicted probabilities between approximately 0.45 and 0.65, producing a smooth probability continuum with weak class separation (**Figure S2A**). This probability plateau resulted in overlap between stable and unstable samples in probability space, introducing ambiguity near the feasibility threshold. Such behavior is undesirable for constrained optimization, as the experimentally observed phase separation outcome is strictly binary, with no qualitative overlap between stable and unstable formulations.

Parity plots further illustrate this behavior: MLP predictions do not collapse cleanly to 0 or 1 but instead exhibit continuous variation across the probability axis (**Figure S2A**). Histogram analysis (**Figure S2A**, accompanying distributions) confirms that a substantial fraction of stable formulations occupies mid-range probability bins, outside of confident classification.

In contrast, the RF classifier produced two sharply separated probability clusters. Stable formulations were assigned tightly grouped low probabilities (approximately 0.10–0.15), while unstable formulations formed a compact high-probability cluster (approximately 0.75–0.80). The corresponding parity plot exhibits near-binary behavior with minimal overlap between classes (**Figure S2A**). This low-entropy probability structure is advantageous for enforcing feasibility constraints in Bayesian optimization.

***Thresholded Classification Performance***

Confusion matrix analyses at a fixed probability threshold of 0.5 further reinforced the differences between the two classifiers (**Figure S2B**). The RF classifier achieved perfect classification performance, with zero false positives and zero false negatives across all folds. In contrast, the MLP classifier misclassified several stable formulations as unstable (five false positives), consistent with its mid-range probability plateau.

In a constrained optimization context, such false positives would lead to unnecessary rejection of experimentally viable formulations, artificially restricting the feasible design space. The absence of false positives in the RF classifier provides a practical advantage when feasibility probabilities are used directly within the acquisition function.

Taken together, these diagnostic analyses support the selection of the calibrated random forest classifier as the Phase-1 feasibility surrogate for constrained Bayesian optimization.

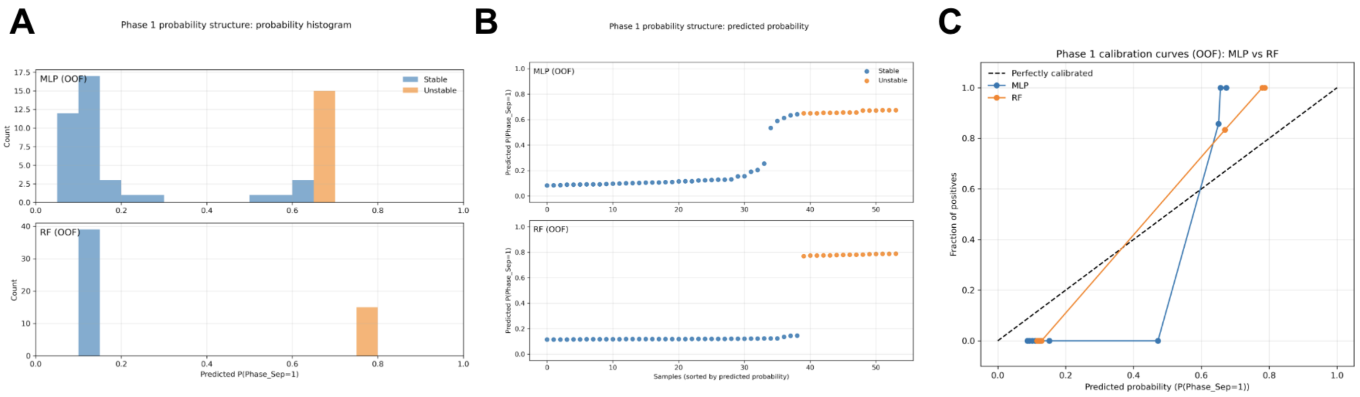


***Figure S3.*** *Additional Phase 1 Modeling Results.* **(A)** Phase 1 probability histograms (MLP vs RF). Histograms of out-of-fold predicted instability probabilities for stable and unstable formulations. The MLP (top) exhibits broader probability spread for stable samples, while the RF (bottom) produces highly concentrated probability distributions, reflecting stronger class certainty. **(B)** Phase 1 predicted probability structure (MLP vs RF). Out-of-fold predicted probabilities for Phase 1 classification, sorted by predicted instability probability. Results are shown for the multilayer perceptron (MLP, top) and random forest (RF, bottom). Stable and unstable formulations separate into distinct probability regions, illustrating model confidence and class separation. **(C)**. Phase 1 calibration curves (MLP vs RF). Out-of-fold calibration curves comparing predicted instability probabilities to observed outcome frequencies for MLP and RF classifiers. The dashed line indicates perfect calibration. RF predictions exhibit closer alignment with ideal calibration across the probability range.

***Table S5.*** *Comparison of calibrated Random Forest and MLP classifiers for Phase 1 stability prediction.* Out-of-fold (OOF) performance metrics were calculated using nested five-fold cross-validation with probability calibration applied within each fold.

| Metric | Random Forest (RF) | MLP Classifier |
|---|---|---|
| CV Best Negative Log-Loss | -0.0063 | -0.3402 |
| Calibration Method | Sigmoid | Sigmoid |
| Log-Loss (OOF) | 0.1618 | 0.2803 |
| ROC-AUC | 1.00 | 1.00 |
| PR-AUC | 1.00 | 1.00 |
| Brier Score | 0.0239 | 0.0752 |
| Accuracy | 1.00 | 0.907 |
| F1 | 1.00 | 0.857 |
| Mean Predictive Entropy | 0.412 | 0.464 |

### ***Phase 2 Supplemental Information***

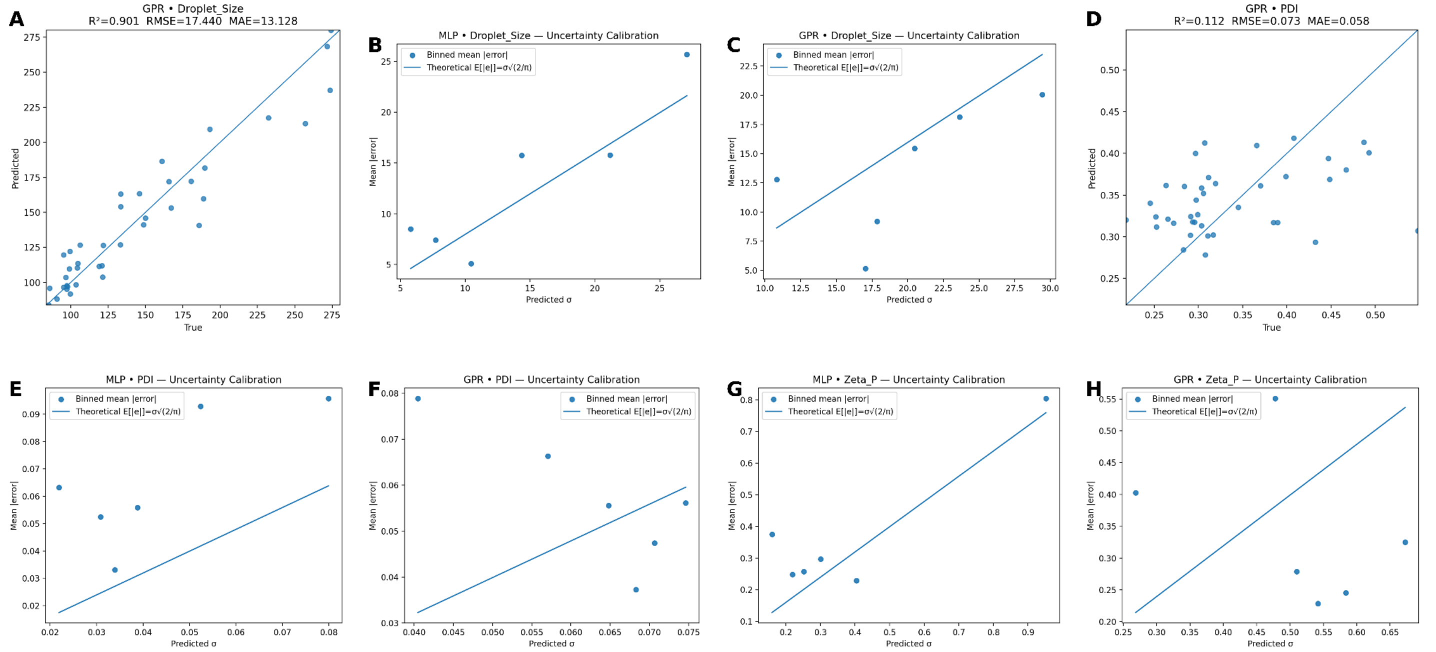


***Figure S4****. Phase 2 Surrogate Model Performance and Calibration.* GPR parity plots, MLP uncertainty calibrations, and GPR uncertainty calibrations are shown for (**A-C)** Droplet Size and **(D-F**) PDI, respectively. (**G**) and (**H**) show the MLP and GPR uncertainty calibrations for zeta potential.

##### ***Uncertainty Calibration Diagnostics***

Uncertainty calibration behavior was evaluated explicitly using parity plots and uncertainty calibration curves (**Figures S3-4**). These diagnostics were treated as first-class selection criteria alongside likelihood metrics.

For droplet size, the MLP ensemble exhibited overconfidence across multiple uncertainty regimes, with empirical errors exceeding theoretical expectations for a calibrated Gaussian predictor (**Figure S4B**). In contrast, GPR uncertainty estimates were generally conservative, with empirical errors at or below theoretical expectations across bins (**Figure S4C**), motivating the selection of GPR despite marginally higher NLL.

For PDI, GPR uncertainty estimates closely tracked empirical residuals across uncertainty bins (**Figure S2F**), while MLP exhibited pronounced overconfidence in the low-σ regime (**Figure S2E**), reinforcing the selection of GPR for this target.

For zeta potential, the trend was reversed. GPR exhibited systematic underconfidence, with predictive uncertainties exceeding observed errors (**Figure S4H**). The MLP ensemble produced uncertainty behavior more closely aligned with theoretical expectations (**Figure S3G**), supporting its selection for this response.

***Table S6.*** *Phase 2 regression surrogate performance and NLL-based rankings.*
Model performance for physically stable formulations (n = 39) was evaluated using nested five-fold cross-validation. Reported metrics are out-of-fold root mean squared error (RMSE), mean absolute error (MAE), coefficient of determination ($R^2$), and negative log-likelihood (NLL). Random Forest (RF) and Support Vector Regression (SVR) models serve as benchmark baselines. Rankings are computed within each target variable, with rank 1 indicating the best performance. Bold rows indicate surrogates selected for deployment in Bayesian optimization.

| Model | OOF RMSE | OOF MAE | OOF $R^2$ | OOF NLL | rank NLL | rank RMSE | rank $R^2$ |
|---|---|---|---|---|---|---|---|
| Average Droplet Size | | | | | | | |
| MLP | 16.89 | 12.57 | 0.91 | 4.10 | 1 | 1 | 1 |
| GPR | 17.44 | 13.13 | 0.90 | 4.41 | 2 | 2 | 2 |
| Polydispersity Index | | | | | | | |
| MLP | 0.08 | 0.06 | -0.17 | 0.58 | 2 | 2 | 2 |
| GPR | 0.07 | 0.06 | 0.11 | -0.91 | 1 | 1 | 1 |
| Zeta Potential | | | | | | | |
| MLP | 0.52 | 0.36 | 0.69 | 0.87 | 1 | 2 | 2 |
| GPR | 0.45 | 0.34 | 0.77 | 0.94 | 2 | 1 | 1 |

##### *Metric-Based Screening of Candidate Surrogates*

Out-of-fold regression performance metrics for all candidate Phase-2 surrogate models are summarized in Table S4. Models were evaluated using RMSE, MAE, coefficient of determination ($R^2$), and negative log-likelihood (NLL) under nested five-fold cross-validation. Performance varied substantially across targets, precluding the use of a single global surrogate.

***Droplet size.*** Both MLP ensembles and GPR achieved strong predictive performance, ranking first and second across RMSE, $R^2$, and NLL. The MLP ensemble achieved the lowest NLL, while GPR exhibited comparable likelihood performance and competitive point-estimate accuracy.

***Polydispersity index (PDI).*** PDI proved the most challenging target, with $R^2$ values ranging from −0.3 to 0.2 across models. GPR outperformed other candidates, achieving the lowest NLL, competitive RMSE and MAE, and a positive coefficient of determination.

***Zeta potential.*** The MLP ensemble achieved the strongest overall performance, ranking favorably across RMSE, $R^2$, and NLL. GPR underperformed relative to MLP in both point accuracy and probabilistic scoring.

Although metric-based screening narrowed the candidate set primarily to GPR and MLP models, additional diagnostic evaluation was required to assess uncertainty suitability for Bayesian optimization.

#### ***Modeling Approach and Results Supplemental Information***

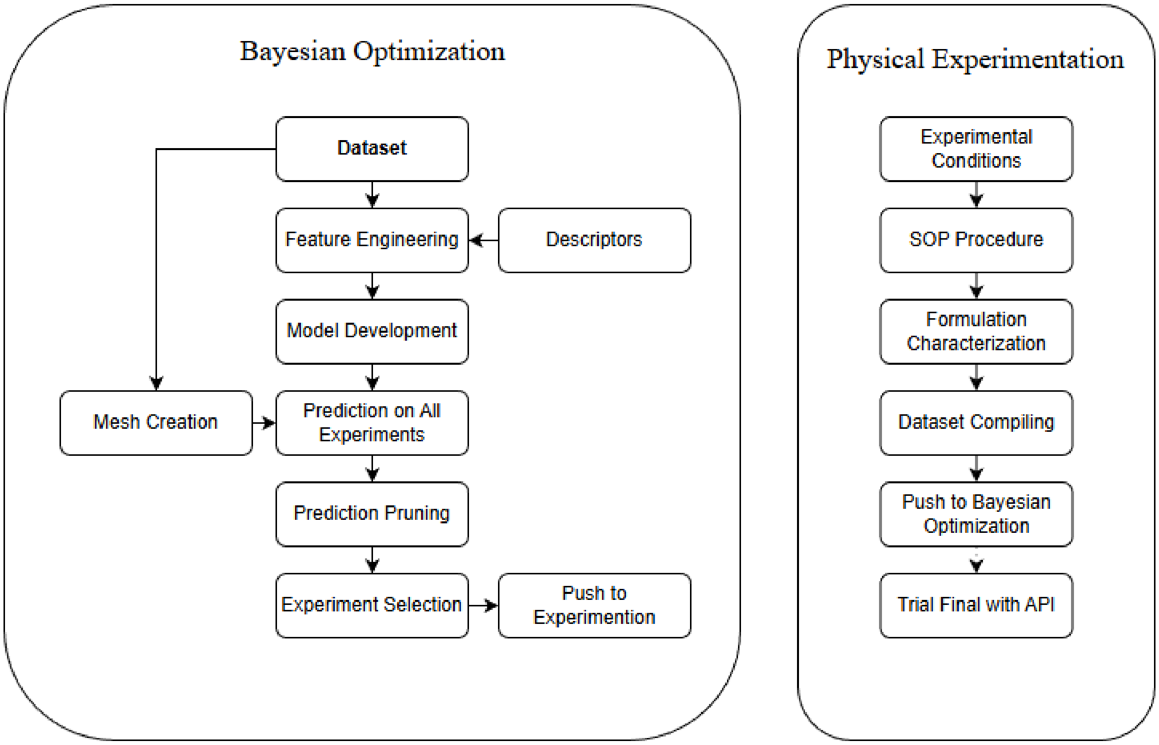


***Figure S5****. General Machine Learning Campaign Process Flow Diagram.* Overview of the experimentation and modeling workflow. The cycle begins at the Dataset node in Bayesian Optimization. Optimization algorithm is developed, used to predict upon all possible experiments, then the 5 best performing microemulsions from the prediction set are selected to move to the physical experimentation phase. After formulation, the microemulsions are physicochemically characterized and their data is tabulated and added back to the working dataset. The cycle is completed when the maximum experimental burden is reached.

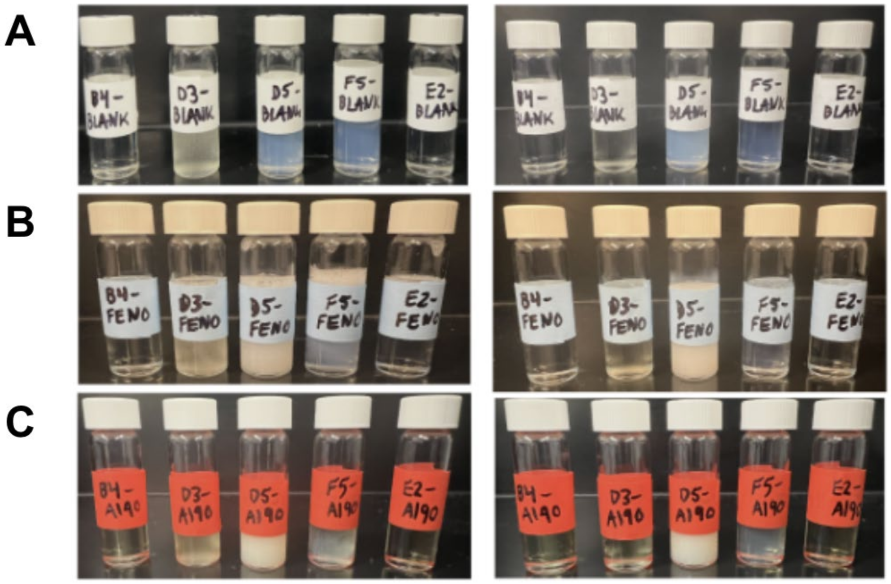


***Figure S6****. Microemulsions during early phase of stability study*. Microemulsions are shown immediately after formulation (left-hand panels) and after 24 hours of storage (right-hand panels) prior to characterization. (**A**) Blank, (**B**) Fenofibrate-, and (**C**) A190-loaded formulations are shown.

***Table S7.*** *Physicochemical Outcomes for Top-Performing Microemulsions.*
Values displayed are those resultants from the optimization campaign, and were measured after 24 hr storage at room temperature. Data are presented as Mean ± SD (n=3).

| Formulation | Average Droplet Size (nm) | Polydispersity Index | Zeta Potential (mV) |
|---|---|---|---|
| B4 | 11.69 ± 1.05 | 0.170 ± 0.081 | -3.40 ± 0.27 |
| D3 | 15.28 ± 1.94 | 0.186 ± 0.066 | -2.23 ± 0.24 |
| D5 | 132.21 ± 1.22 | 0.296 ± 0.001 | -3.07 ± 0.63 |
| F5 | 148.14 ± 2.11 | 0.269 ± 0.015 | -3.01 ± 0.68 |
| E2 | 12.40 ± 2.24 | 0.216 ± 0.093 | -4.88 ± 1.46 |

***Table S8.*** *Physicochemical Outcomes of Reformulated, Drug-Loaded Microemulsions*
Values displayed are those resultant from the modeling campaign and the post-campaign stability study described. Formulations were characterized after 24 hr storage at room temperature. Data are presented as Mean ± SD (n=3).

| Formulation | Average Droplet Size (nm) | | Polydispersity Index | | Zeta Potential (mV) | |
|---|---|---|---|---|---|---|
| | Campaign | Post-Campaign | Campaign | Post-Campaign | Campaign | Post-Campaign |
| B4 | 11.69 ± 1.05 | 151.4 ± 1.4 | 0.170 ± 0.081 | 0.229 ± 0.011 | -3.40 ± 0.27 | -1.15 ± 0.30 |
| D3 | 15.28 ± 1.94 | 78.01 ± 2.02 | 0.186 ± 0.066 | 0.703 ± 0.012 | -2.23 ± 0.24 | -2.22 ± 0.16 |
| D5 | 132.21 ± 1.22 | 115.87 ± 1.37 | 0.296 ± 0.001 | 0.104 ± 0.009 | -3.07 ± 0.63 | -2.64 ± 0.07 |
| F5 | 148.14 ± 2.11 | 158.53 ± 6.12 | 0.269 ± 0.015 | 0.296 ± 0.021 | -3.01 ± 0.68 | -2.88 ± 0.25 |
| E2 | 12.40 ± 2.24 | 10.58 ± 0.13 | 0.216 ± 0.093 | 0.168 ± 0.004 | -4.88 ± 1.46 | -3.17 ± 0.22 |

***Table S9***. *Physicochemical Properties of Model Drugs.*

| | A190 | Fenofibrate |
|---|---|---|
| **Chemical Structure** | 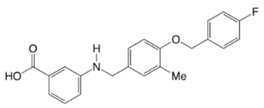 | 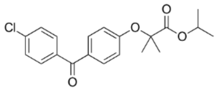 |
| **Molecular Weight** | 365.4 g/mol | 360.8 g/mol |
| **LogP** | 4.8 | 5.2 |
| **Aqueous Solubility** | 0.028 ± 0.01 mg/mL | 0.604 ± 0.002 µg/mL |

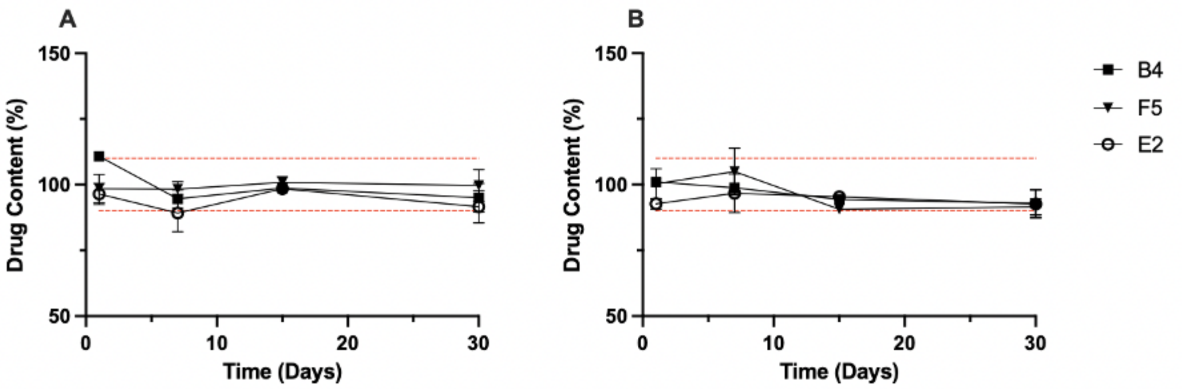


***Figure S7.*** *Drug content of model-optimized microemulsions during 30 days of storage at room temperature*. **(A)** A190-loaded microemulsions and **(B)** Fenofibrate-loaded microemulsions are expressed as percent of initial drug content (%). Data are presented as Mean ± SD (n=3), and the red dashed lines indicate 90 to 110% of the initial drug content for the respective formulation.

***Table S10.*** *Statistical Analysis of PAMPA Permeability Results*

| | ANOVA [a] | Post-Hoc Testing [b] | | |
|---|---|---|---|---|
| **Drug** | **p-value** | **Comparison** | **Mean Difference** | **p-value** |
| A190 | <0.001 | B4 vs F5 | -7.585 ± 0.9509 | < 0.001 |
| | | F5 vs E2 | 0.8590 ± 1.009 | 0.911 |
| | | B4 vs E2 | -6.726 ± 0.9104 | < 0.001 |
| Fenofibrate | <0.001 | B4 vs F5 | -5.369 ± 0.8051 | < 0.001 |
| | | F5 vs E2 | -1.983 ± 0.8540 | 0.186 |
| | | B4 vs E2 | -7.352 ± 0.8540 | <0.001 |

a. An Ordinary One-way ANOVA was used to evaluate differences among groups;
b. Post-hoc analyses were performed using Tukey's HSD. Values are presented as Mean ± SEM (n=4-6). A p-value < 0.05 was considered statistically different.

This is the end of the supplemental figures and tables, which are referenced in the main manuscript. The following are additional figures that give further insight into batching, data acquisition, and experimental space explored within the optimization campaign.

### ***Acquisition and Objective Function Supplemental Information***

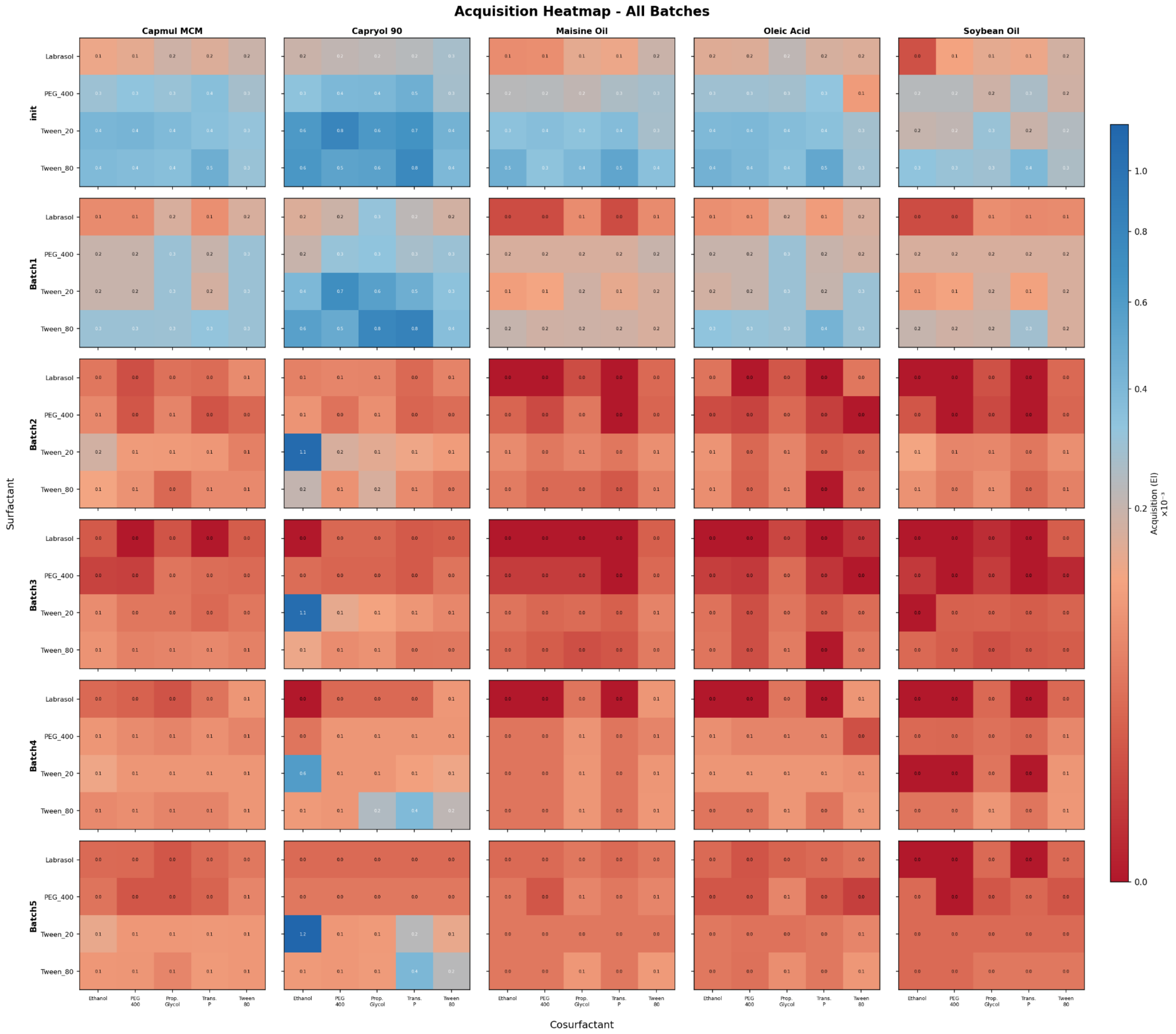


***Figure S8.*** *Acquisition Function Heat Map on Categorical Combinations.* Heatmap visualization of the acquisition function on the first experiment of each batch from the initial dataset (init) to potential Batch 6 acquisition, and the evolution of which categorical combinations are most beneficial to the Bayesian optimization campaign. Init represents the acquisition for the first experiment for batch 1.

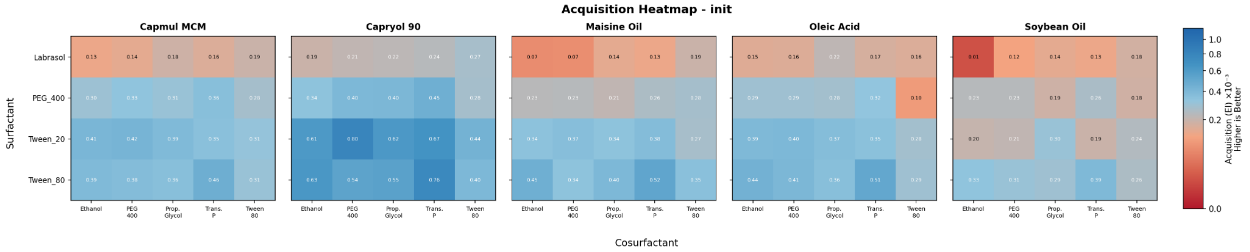


***Figure S9.*** *Acquisition Function Heat Map on Categorical Combination of Initial Dataset for Batch 1.* Zoom on Batch 1's acquisition function values for experiment 1 of the batch using the initial dataset.

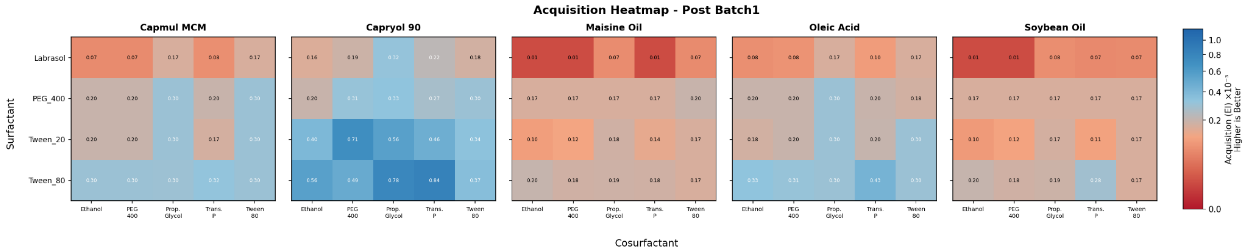


***Figure S10.*** *Acquisition Function Heat Map on Categorical Combination of Intermediate Dataset 1 for Batch 2.* Zoom on Batch 2's acquisition function values for experiment 1 of the batch using the intermediate dataset 1, consisting of the initial dataset with experimental results from batch 1.

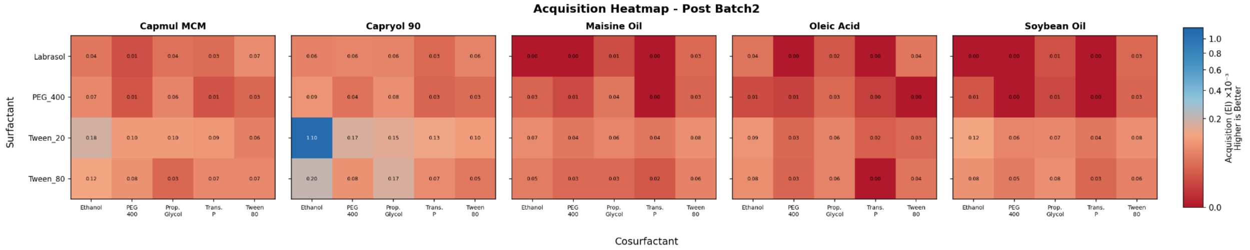


***Figure S11.*** *Acquisition Function Heat Map on Categorical Combination of Intermediate Dataset 2 for Batch 3* Zoom on Batch 3's acquisition function values for experiment 1 of the batch using the intermediate dataset 2, consisting of the initial dataset with experimental results from batch 1-2.

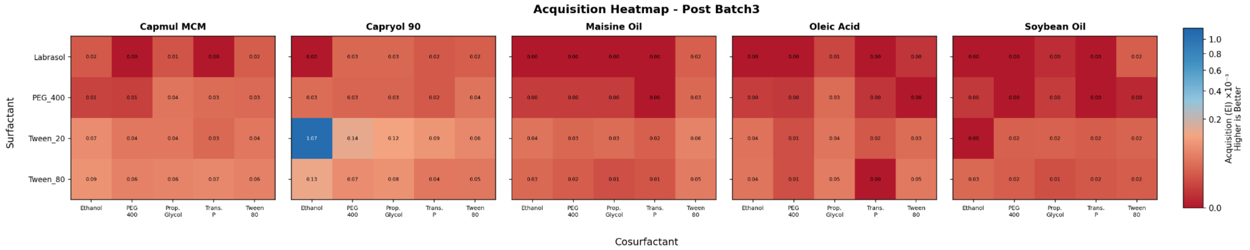


***Figure S12.*** *Acquisition Function Heat Map on Categorical Combination of Intermediate Dataset 3 for Batch 4* Zoom on Batch 4's acquisition function values for experiment 1 of the batch using the intermediate dataset 3, consisting of the initial dataset with experimental results from batch 1-3.

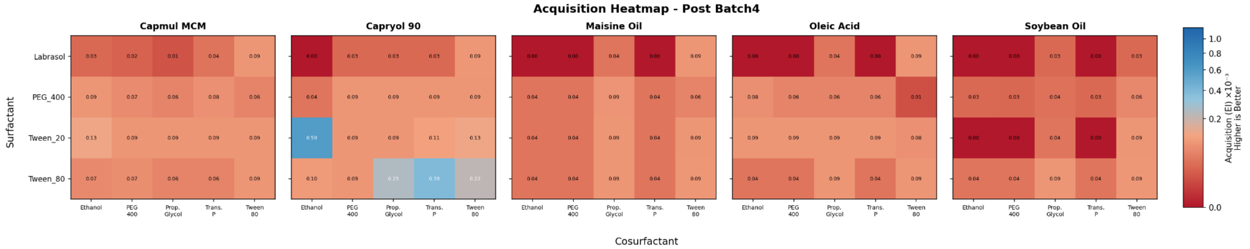


***Figure S13.*** *Acquisition Function Heat Map on Categorical Combination of Intermediate Dataset 4 for Batch 5.* Zoom on Batch 5's acquisition function values for experiment 1 of the batch using the intermediate dataset 4, consisting of the initial dataset with experimental results from batch 1-4.

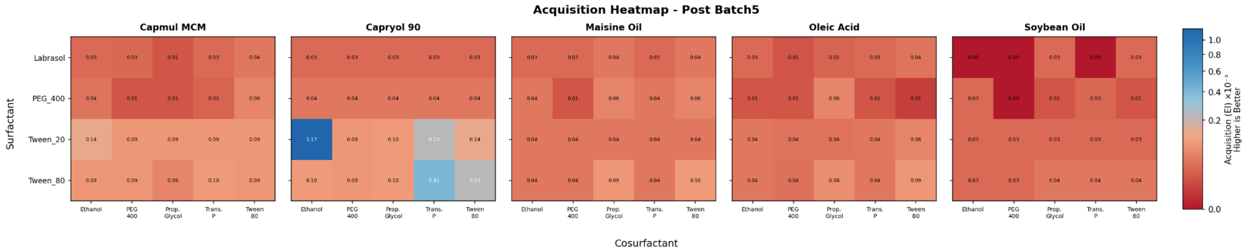


***Figure S14.*** *Acquisition Function Heat Map on Categorical Combination of Final Dataset for Batch 6.* Zoom on potential Batch 6's acquisition function values for experiment 1 of the batch using the final dataset, consisting of the initial dataset with experimental results from batch 1-5.

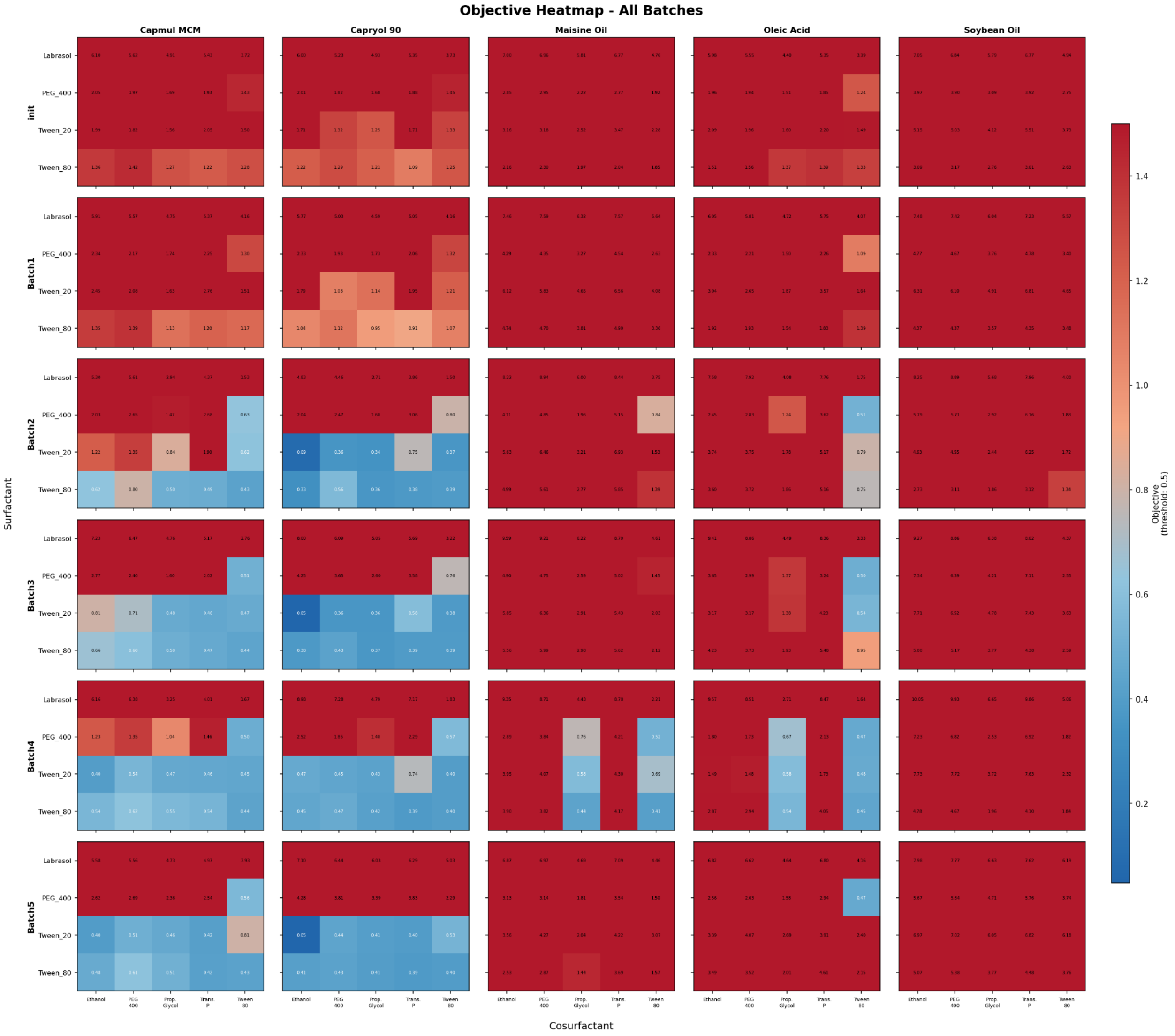


***Figure S15.*** *Objective Function Heat Map on Categorical Combinations.* Heatmap visualization of the objective function on the first experiment of each batch from the initial dataset (init) to potential Batch 6 objective, and the evolution of which categorical combinations areas are most predicted to perform the best by the multiphase surrogate model. Init represents the acquisition for the first experiment for batch 1.

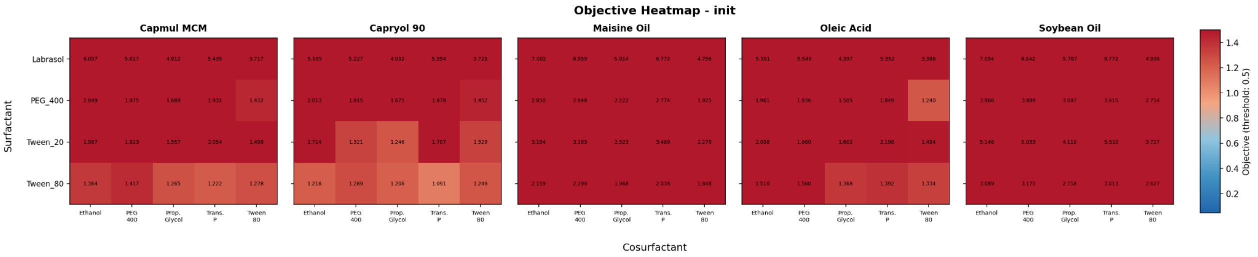


***Figure S16.*** *Objective Function Heat Map on Categorical Combination of Initial Dataset for Batch 1.* Zoom on Batch 1's objective function values for experiment 1 of the batch using the initial dataset.

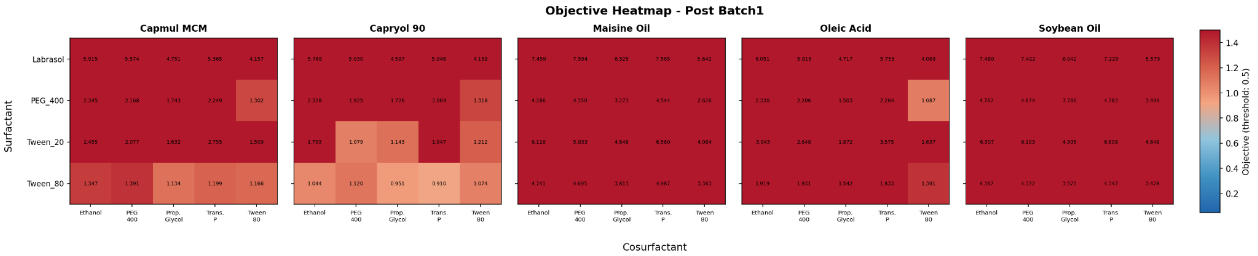


***Figure S17.*** *Objective Function Heat Map on Categorical Combination of Intermediate Dataset 1 for Batch 2.* Zoom on Batch 2's objective function values for experiment 1 of the batch using the intermediate dataset 1, consisting of the initial dataset with experimental results from batch 1.

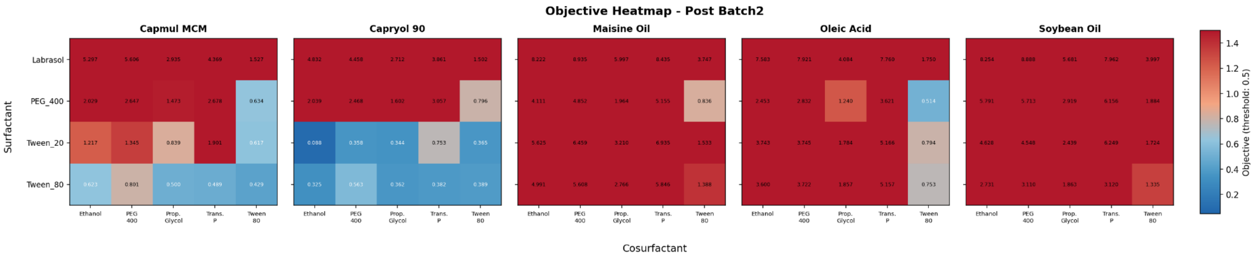


***Figure S18.*** *Objective Function Heat Map on Categorical Combination of Intermediate Dataset 2 for Batch 3.* Zoom on Batch 3's objective function values for experiment 1 of the batch using the intermediate dataset 2, consisting of the initial dataset with experimental results from batch 1-2.

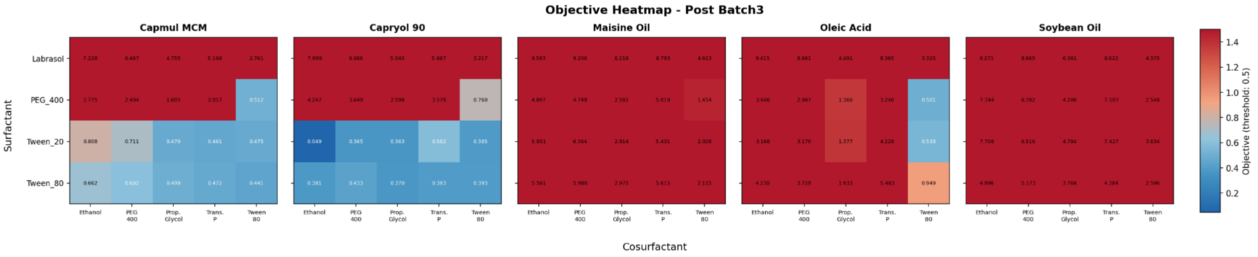


***Figure S19.*** *Objective Function Heat Map on Categorical Combination of Intermediate Dataset 3 for Batch 4.* Zoom on Batch 4's objective function values for experiment 1 of the batch using the intermediate dataset 3, consisting of the initial dataset with experimental results from batch 1-3.

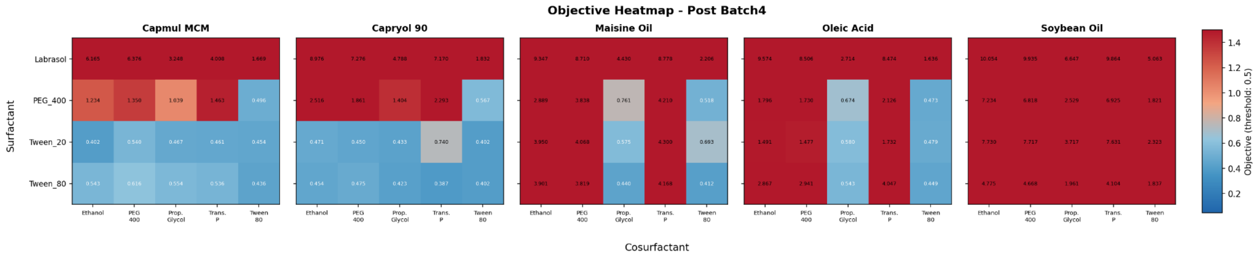


***Figure S20.*** *Objective Function Heat Map on Categorical Combination of Intermediate Dataset 4 for Batch 5.* Zoom on Batch 5's objective function values for experiment 1 of the batch using the intermediate dataset 4, consisting of the initial dataset with experimental results from batch 1-4.

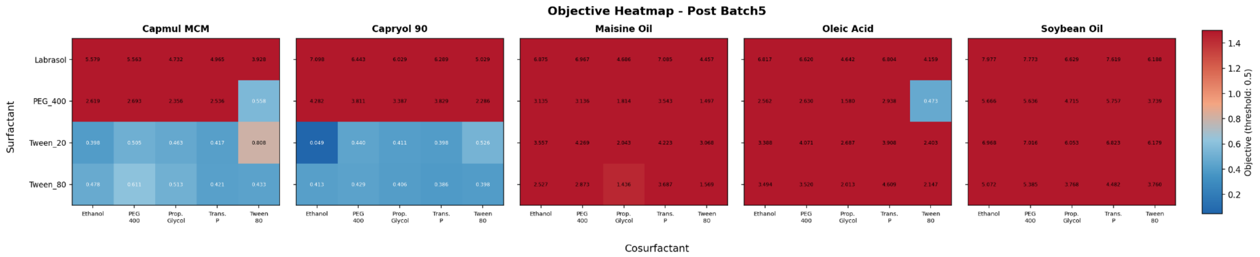


***Figure S21.*** *Objective Function Heat Map on Categorical Combination of Final Dataset for Batch 6.* Zoom on potential Batch 6's objective function values for experiment 1 of the batch using the final dataset, consisting of the initial dataset with experimental results from batch 1-5.

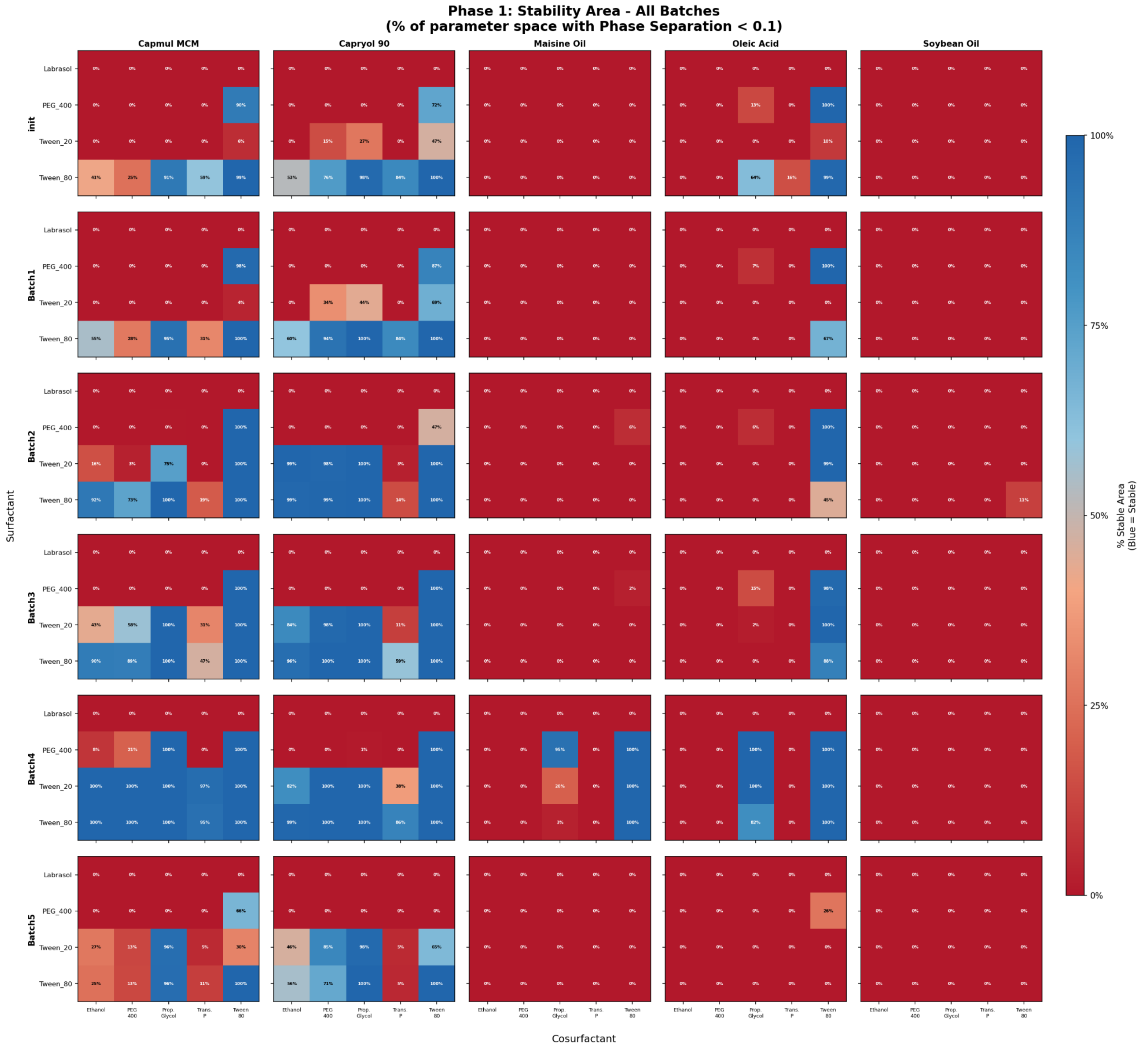


***Figure S22.*** *Phase 1 Surrogate Model Stability Prediction for Every Coarse Mesh.* Heatmap visualization of the predicted stability of each categorical combination at each iteration of the optimization campaign from initial data (init) to final batch (Batch 5). Displays the evolution of the classification model on the coarse mesh, as well as the narrowing down of the design space to a much smaller section of stable formulations to test. The numbers visualized indicate the percentage of the coarse mesh that was identified as stable.

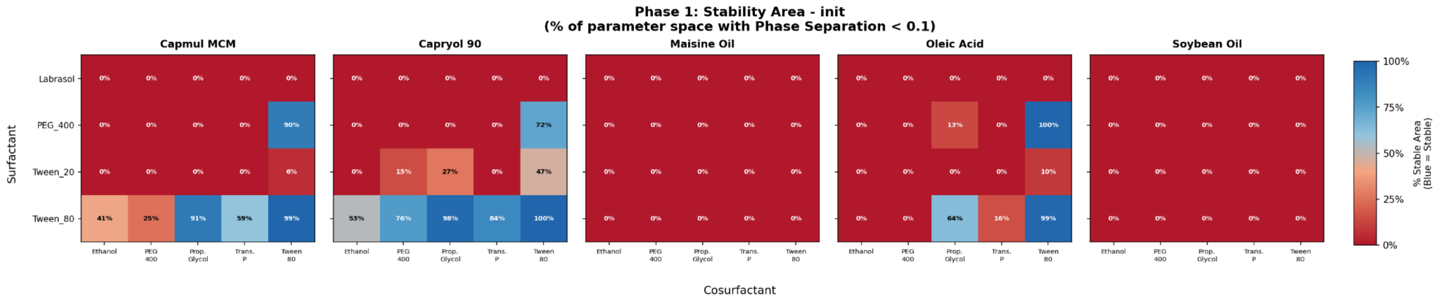


***Figure S23.*** *Phase 1 Surrogate Model Stability Prediction for Batch 1 Coarse Mesh.* Zoom in on the coarse mesh prediction for the first experiment of batch 1 using the initial dataset consisting of design of experiment, miscellaneous, and randomizer datasets.

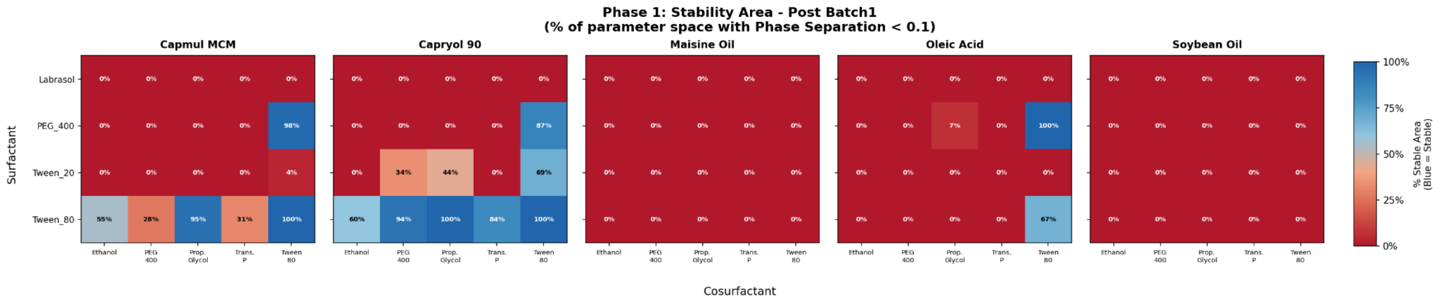


***Figure S24.*** *Phase 1 Surrogate Model Stability Prediction for Batch 2 Coarse Mesh.* Zoom in on the coarse mesh prediction for the first experiment of batch 2 using an intermediate dataset consisting of design of experiment, miscellaneous, randomizer, and batch 1 datasets.

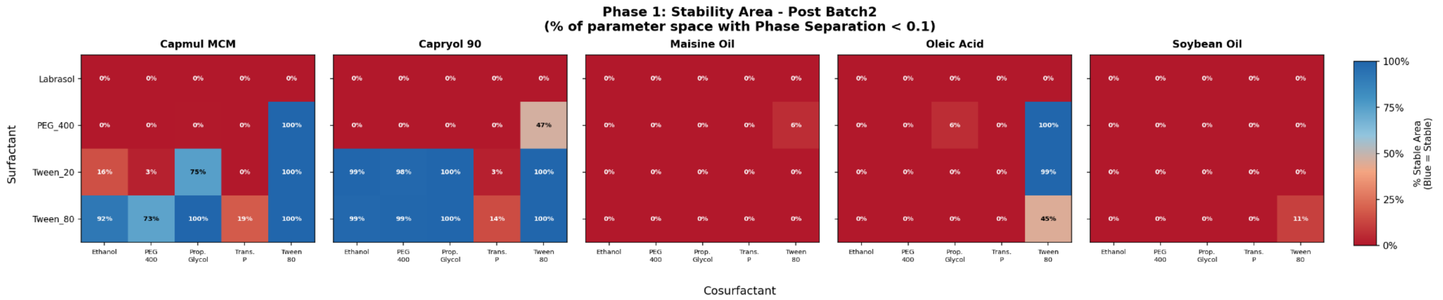


***Figure S25.*** *Phase 1 Surrogate Model Stability Prediction for Batch 3 Coarse Mesh.* Zoom in on the coarse mesh prediction for the first experiment of batch 3 using an intermediate dataset consisting of design of experiment, miscellaneous, randomizer, and batch 1-2 datasets.

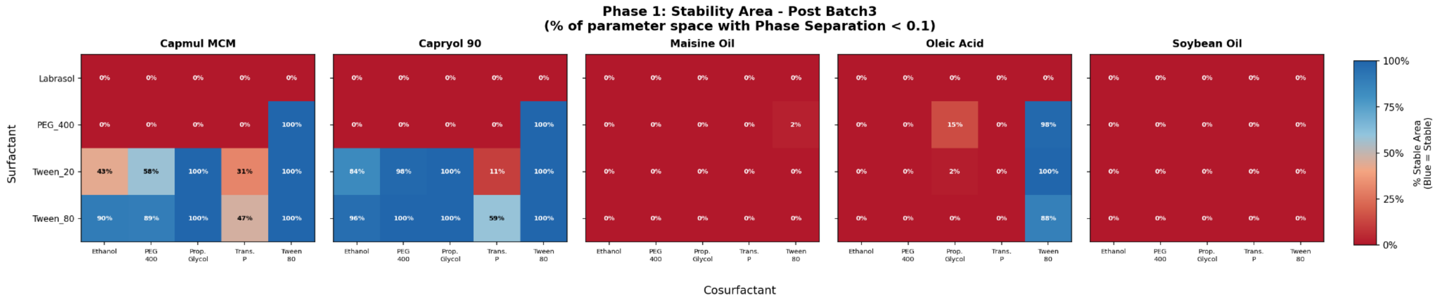


***Figure S26.*** *Phase 1 Surrogate Model Stability Prediction for Batch 4 Coarse Mesh.* Zoom in on the coarse mesh prediction for the first experiment of batch 4 using an intermediate dataset consisting of design of experiment, miscellaneous, randomizer, and batch 1-3 datasets.

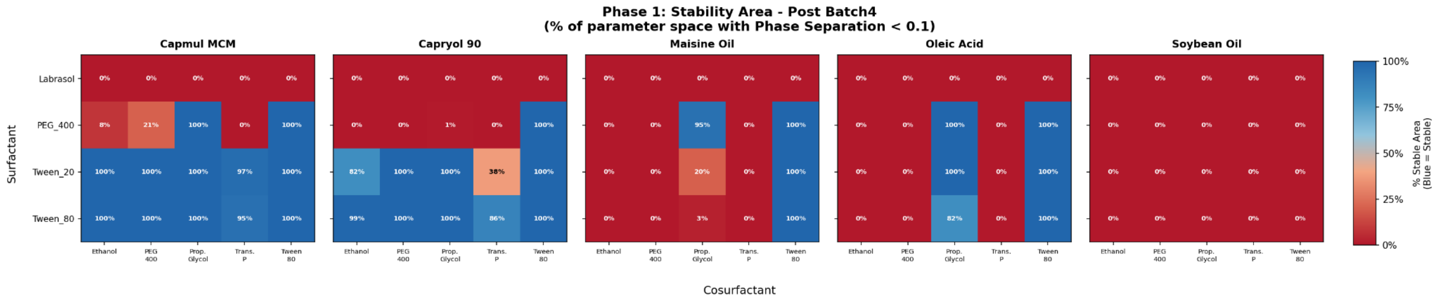


***Figure S27.*** *Phase 1 Surrogate Model Stability Prediction for Batch 5 Coarse Mesh.* Zoom in on the coarse mesh prediction for the first experiment of batch 5 using an intermediate dataset consisting of design of experiment, miscellaneous, randomizer, and batch 1-4 datasets.

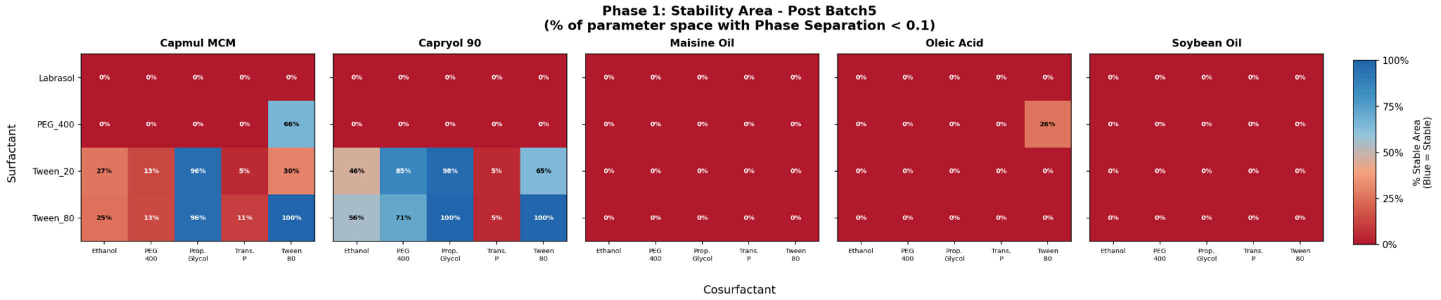


***Figure S28.*** *Phase 1 Surrogate Model Stability Prediction for Potential Batch 6 Coarse Mesh.* Zoom in on the coarse mesh prediction for the first experiment of potential batch 6 using the final dataset consisting of design of experiment, miscellaneous, randomizer, and batch 1-5 datasets.

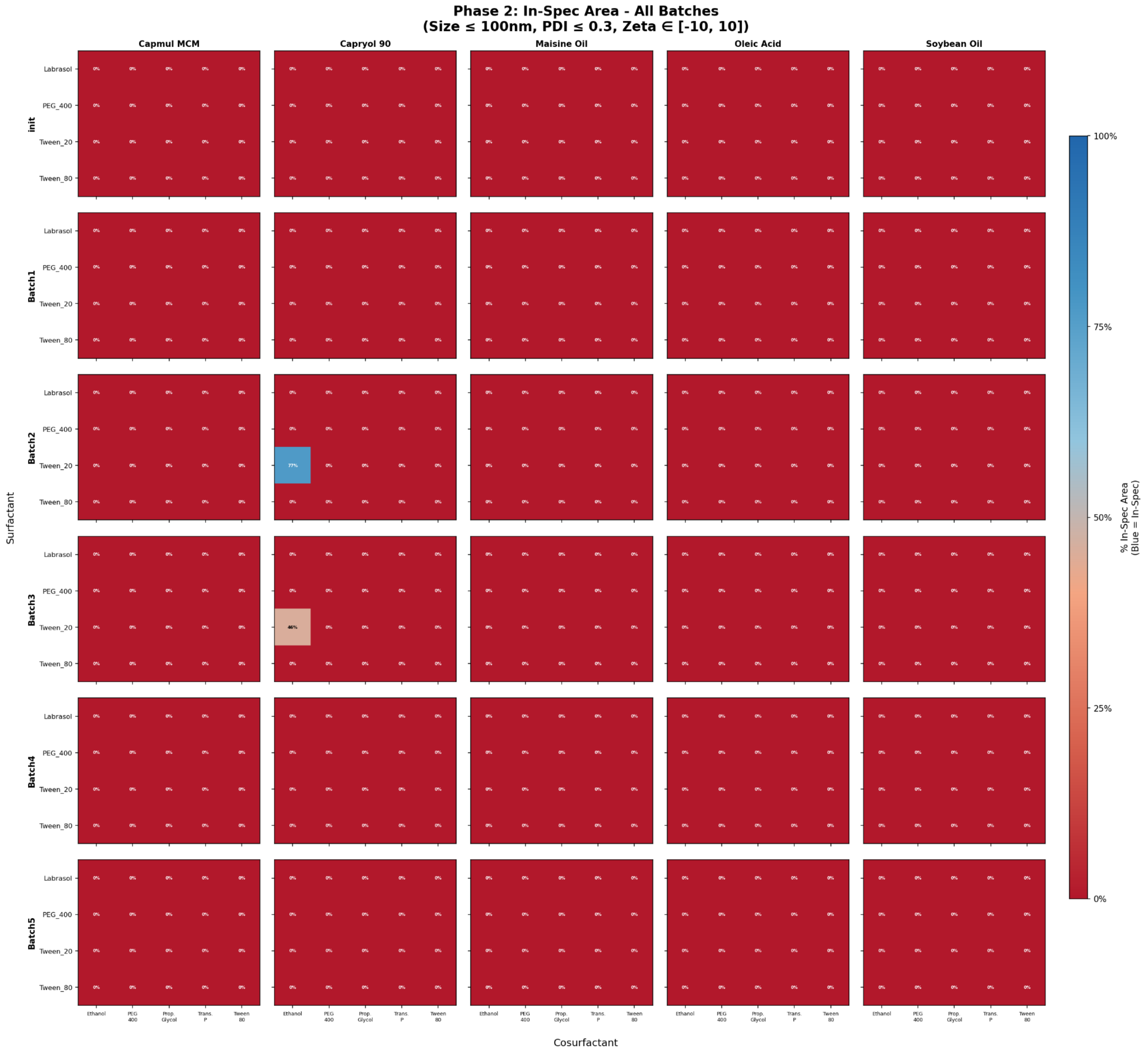


***Figure S29.*** *Phase 2 Surrogate Model Continuous Parameter Prediction for All Coarse Meshes.* Heatmap visualization of the predicted continuous parameters utilizing phase 2 surrogate models for all the coarse meshes at every batch iteration. Displays the evolution of the phase 2 surrogate regression models through each iteration. The heatmap shows how confident the model becomes after obtaining B4 and the regression back to uncertain predictions after obtaining additional data.

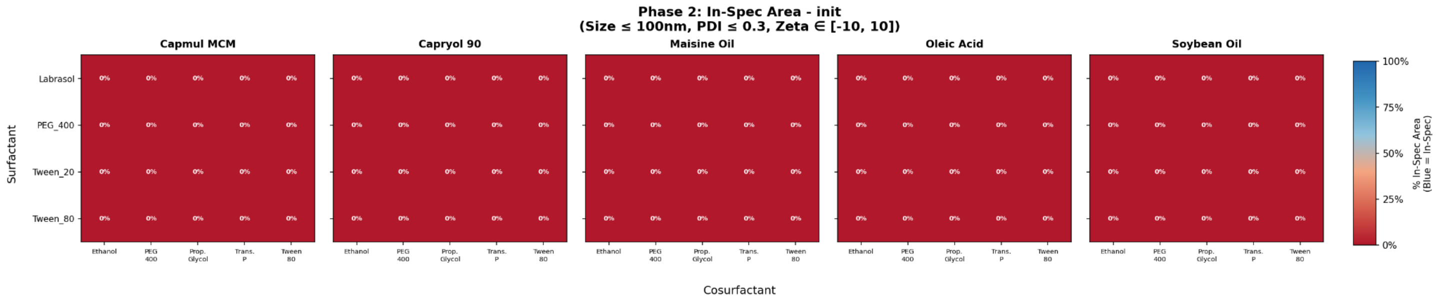


***Figure S30.*** *Phase 2 Surrogate Model Continuous Parameter Prediction for Batch 1 Coarse Mesh.* Zoom in on the coarse mesh prediction for the first experiment of batch 1 using the initial dataset consisting of design of experiment, miscellaneous, and randomizer datasets.

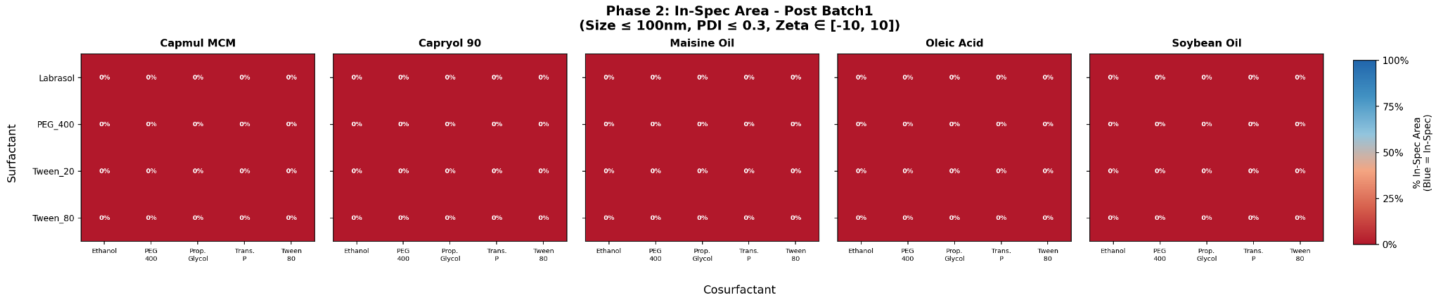


***Figure S31.*** *Phase 2 Surrogate Model Continuous Parameter Prediction for Batch 2 Coarse Mesh.* Zoom in on the coarse mesh prediction for the first experiment of batch 2 using an intermediate dataset consisting of design of experiment, miscellaneous, randomizer, and batch 1 datasets.

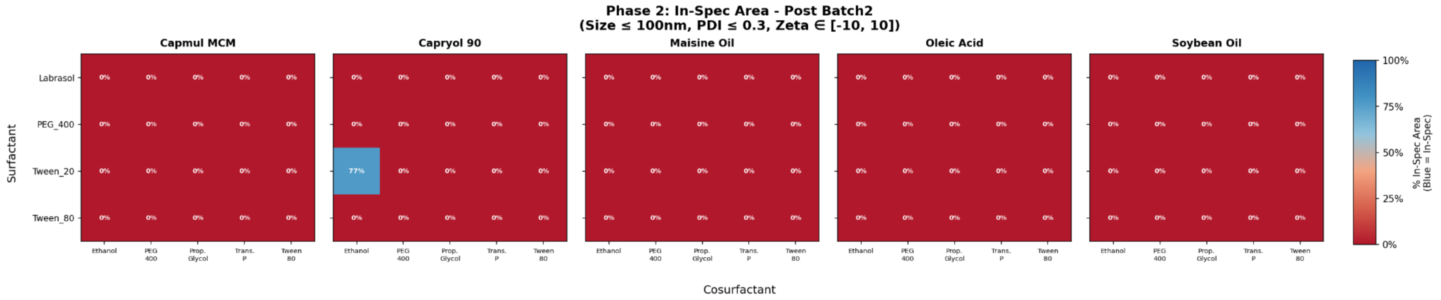


***Figure S32.*** *Phase 2 Surrogate Model Continuous Parameter Prediction for Batch 3 Coarse Mesh.* Zoom in on the coarse mesh prediction for the first experiment of batch 3 using an intermediate dataset consisting of design of experiment, miscellaneous, randomizer, and batch 1-2 datasets.

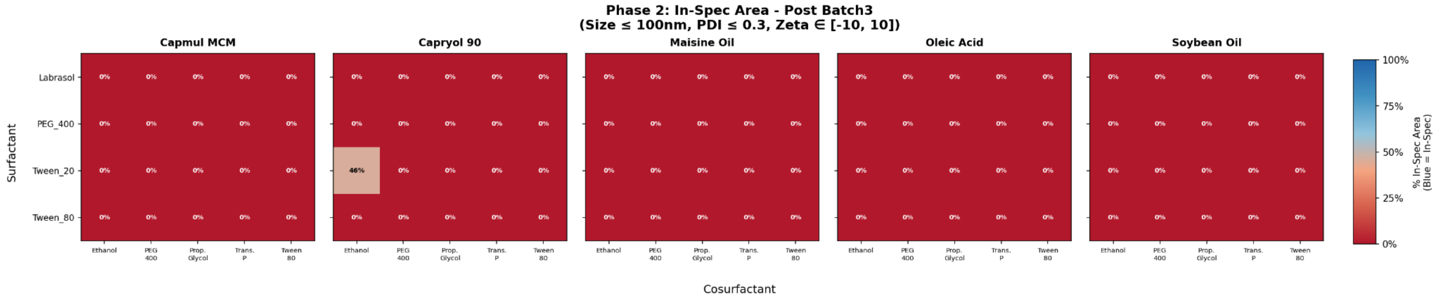


***Figure S33.*** *Phase 2 Surrogate Model Continuous Parameter Prediction for Batch 4 Coarse Mesh.* Zoom in on the coarse mesh prediction for the first experiment of batch 4 using an intermediate dataset consisting of design of experiment, miscellaneous, randomizer, and batch 1-3 datasets.

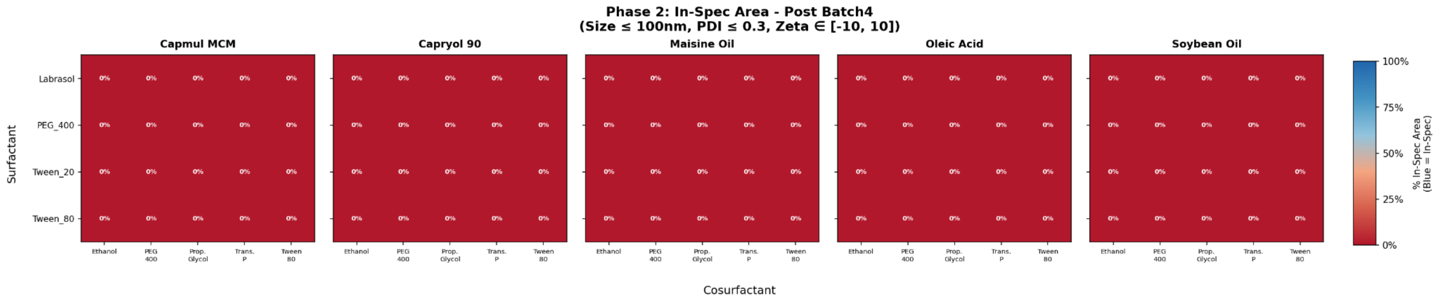


***Figure S34.*** *Phase 2 Surrogate Model Continuous Parameter Prediction for Batch 5 Coarse Mesh.* Zoom in on the coarse mesh prediction for the first experiment of batch 5 using an intermediate dataset consisting of design of experiment, miscellaneous, randomizer, and batch 1-4 datasets.

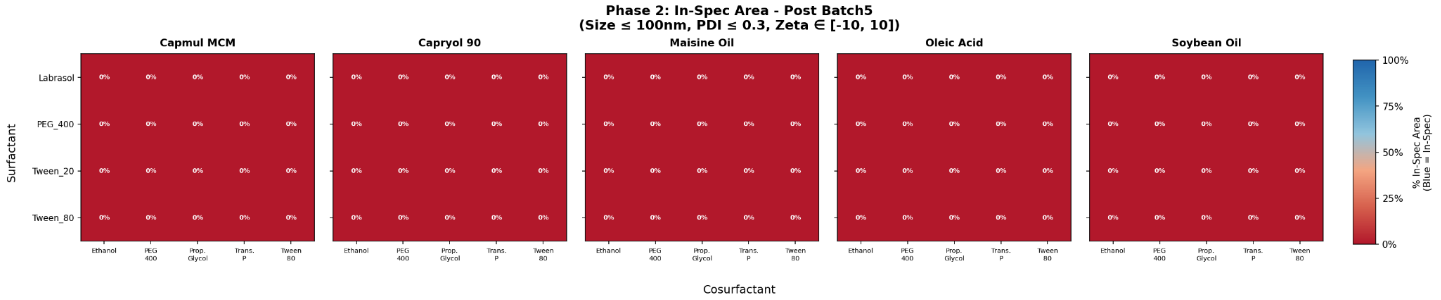


***Figure S35.*** *Phase 2 Surrogate Model Continuous Parameter Prediction for Potential Batch 6 Coarse Mesh.* Zoom in on the coarse mesh prediction for the first experiment of potential batch 6 using the final dataset consisting of design of experiment, miscellaneous, randomizer, and batch 1-5 datasets.

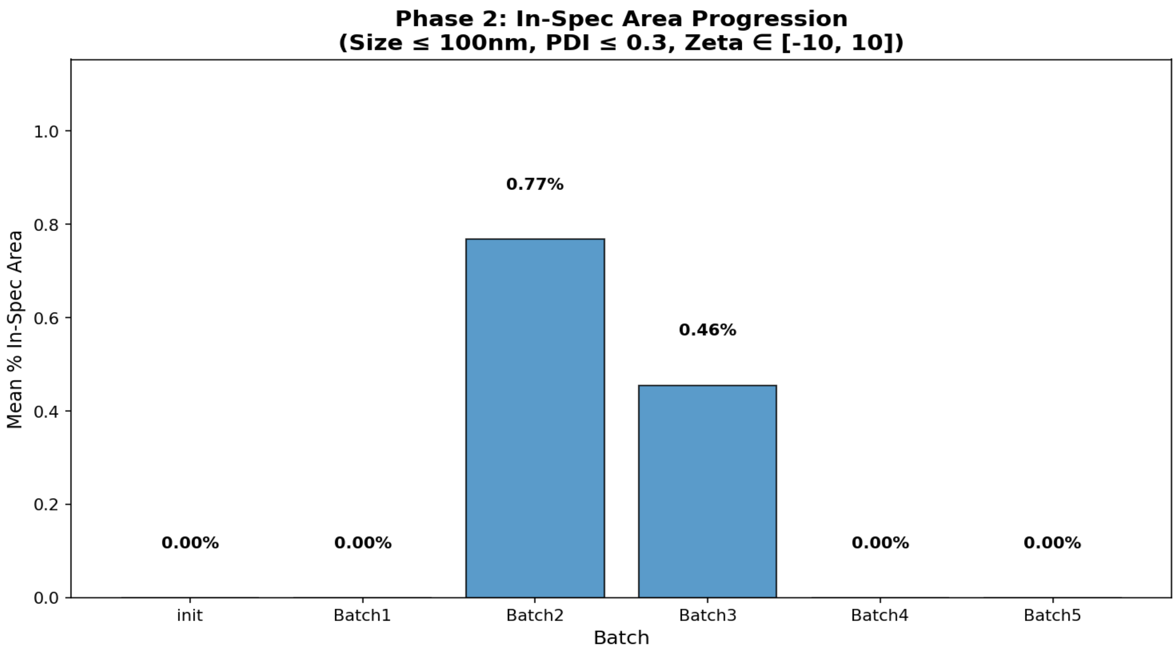


***Figure S36.*** *Phase 2 Surrogate Model Continuous Parameter Prediction In-Specification Percentage.* Evolution of the in-specification percentage of all the coarse meshes given the predication by phase 2 surrogate models.
